# DISCA: high-throughput cryo-ET structural pattern mining by deep unsupervised clustering

**DOI:** 10.1101/2021.05.16.444381

**Authors:** Xiangrui Zeng, Anson Kahng, Liang Xue, Julia Mahamid, Yi-Wei Chang, Min Xu

**Affiliations:** Computational Biology Department, Carnegie Mellon University, Pittsburgh, PA, 15213, USA; Computer Science Department, Carnegie Mellon University, Pittsburgh, PA, 15213, USA; Structural and Computational Biology Unit, European Molecular Biology Laboratory (EMBL), Heidelberg 69117, Germany; Collaboration for joint PhD degree between EMBL and Heidelberg University, Faculty of Biosciences; Department of Biochemistry and Biophysics, Perelman School of Medicine, University of Pennsylvania, Philadelphia, PA, 19104, USA

## Abstract

Cryo-electron tomography directly visualizes heterogeneous macromolecular structures in complex cellular environments, but existing computer-assisted sorting approaches are low-throughput or inherently limited due to their dependency on available templates and manual labels.

We introduce a high-throughput template-and-label-free deep learning approach that automatically discovers subsets of homogeneous structures by learning and modeling 3D structural features and their distributions.

Diverse structures emerging from sorted subsets enable systematic unbiased recognition of macro-molecular complexes *in situ*.

## 1 Main text

In recent years, cryo-Electron Tomography (cryo-ET) has made it possible to image densities of different molecules and their spatial distributions inside intact cells in a near-native, “frozen-hydrated” state to a resolution of a few nanometers in three dimensions [1]. This molecular-resolution visualization of how macromolecular complexes work together inside cells has allowed researchers to obtain mechanistic insights into particular cellular processes and distinguish competing models from one another [2]. However, a major challenge remains to precisely and comprehensively identify densities of different molecules in complex cellular tomograms. A popular method to perform this task is “template matching” [3], which uses available structures obtained *in vitro* from X-ray crystallography, nuclear magnetic resonance spectroscopy, or single-particle cryo-electron microscopy as template references to search for similar shapes in the tomograms. While useful, its dependency on available structural templates may introduce reference-dependent bias [4], especially when a template is only available from a different species than the one imaged. An alternative popular practice is to manually pick target structures and then average them to obtain the initial template, which is also biased by subjective preferences [5]. More importantly, as evidenced by genome sequencing and mass spectrometry, the native structure of a large number of macromolecular complexes remains unknown [6,7]. Macromolecular complexes that lack available structural information cannot be identified in cryo-ET cellular tomograms using existing structural templates.

With that in mind, we and others have previously proposed a structural pattern mining approach [8,9], as an important step towards template-free visual proteomics [10]. This approach consists of (1) template-free particle picking steps that detect potential structures in a tomogram and (2) recognition steps that classify each particle as a particular type of structure. However, the throughput of these methods is limited because they involve a tremendous number of geometric transformation operations. With the recent advance of cryo-ET data collection methods [11,12], large numbers of tomograms can now be produced daily (50 tomograms of size ~4, 000×6, 000×1, 000 voxels, containing up to a million particles), allowing the effective imaging of many samples with different treatments and experimental controls for comparative analyses. The computationally expensive structural pattern mining approaches are impractical for handling such large-scale datasets. A new type of high-throughput analysis method is therefore needed to allow systematic and comprehensive investigation of the fast-growing size of *in situ* cryo-ET data.

Recently, supervised deep learning methods have been gaining momentum for cryo-ET image analysis [15,16]. By automatically learning better heuristics from accumulating data, their accuracy can improve over time, and they have been shown to perform much more efficiently and accurately than the aforementioned traditional geometry-based approaches [16,17]. Due to their significantly faster recognition speed, they also promise better scalability to large-scale datasets with a large number of classes encompassing heterogeneous structures. However, supervised methods pose an additional major challenge: creating valid training data. In all these supervised deep learning methods, training a neural network requires a substantial amount of pre-labeled data. For cryo-ET, training data has conventionally been produced either by using template matching mentioned above or via laborious manual labeling of target structural patterns in tomograms [15]. Both unavoidably produce reference-dependent biases that limit the analysis. Unfortunately, this difficulty cannot be circumvented by using an annotated tomogram database consisting of multiple independent sources as a less-biased universal training set. This difficulty is because training from separated cryo-ET data sources, collected under different imaging conditions, was shown to result in lower recognition accuracy due to the variable image intensity distribution among data sources [18,19]. Moreover, these supervised methods remain unable to discover structures that are not annotated in the training dataset, posing a similar limitation to template matching. Therefore, a more natural and effective approach could be training the neural network in an unbiased template-and-label-free way by using comprehensive intrinsic structural features in the data themselves.

In light of this, we introduce a high-throughput unsupervised learning approach, DISCA (Deep Iterative Subtomogram Clustering Approach). DISCA automatically discovers structurally homogeneous particle subsets in large-scale cryo-ET datasets by learning 3D structural features extracted by a Convolutional Neural Network (CNN) and statistically modeling the feature distributions. Given a dataset of reconstructed 3D tomograms, as a preprocessing step, we first use template-free particle picking to detect potential structures and extract them as subtomograms. The extracted subtomograms contain heterogeneous structures. We then use DISCA (**Fig. 1**) to sort the subtomograms into relatively homogeneous structural subsets. Specifically, we formulate a generalized Expectation-Maximization (EM) framework (**Supplementary Fig. 1** and **2**) that iteratively clusters subtomograms based on their extracted CNN features and optimizes the CNN through unsupervised training. Finally, as postprocessing steps done outside our framework, the sorted subsets are aligned, averaged and re-embedded to the original tomogram space to validate the recovered structures and their spatial distributions. The design of DISCA enables transformation-invariant feature extraction, automatic estimation of the number of clusters, and progressively improved performance with larger sample sizes, which are validated on realistically simulated datasets of various imaging parameters (**Supplementary Note 1**). For feature extraction in DISCA, we designed a special CNN named YOPO (You Only Pool Once) (**Supplementary Fig. 3**). YOPO preserves detailed structural information and extracts rotation- and translation-invariant features from subtomogram data. Such invariance usually cannot be achieved by standard CNN architecture designs. As independently evaluated by the SHape REtrieval Contest (SHREC) 2020 [16] in a supervised learning task, YOPO achieved the third-best accuracy and outperformed the template matching baselines. Most importantly, YOPO only requires localized coordinates of target macromolecules for training, in which, a whole subtomogram only needs a single label. In comparison, all the other participating methods require labeled segmentation maps for training, in which every voxel needs to be labeled. The segmentation maps (dense labels) for a real cryo-ET dataset are extremely time-consuming to prepare as every single voxel of a tomogram needs to be labeled by experts. Therefore, YOPO was deemed ‘significantly more accessible for cryo-ET researchers’ given that a minimal amount of training supervision was needed [16]. We note that, in DISCA, the training of YOPO is completely unsupervised and further automated to be free from all external domain knowledge, including existing structural templates, manual labeling, or manual selection of densities in the tomograms.

**Figure 1:**
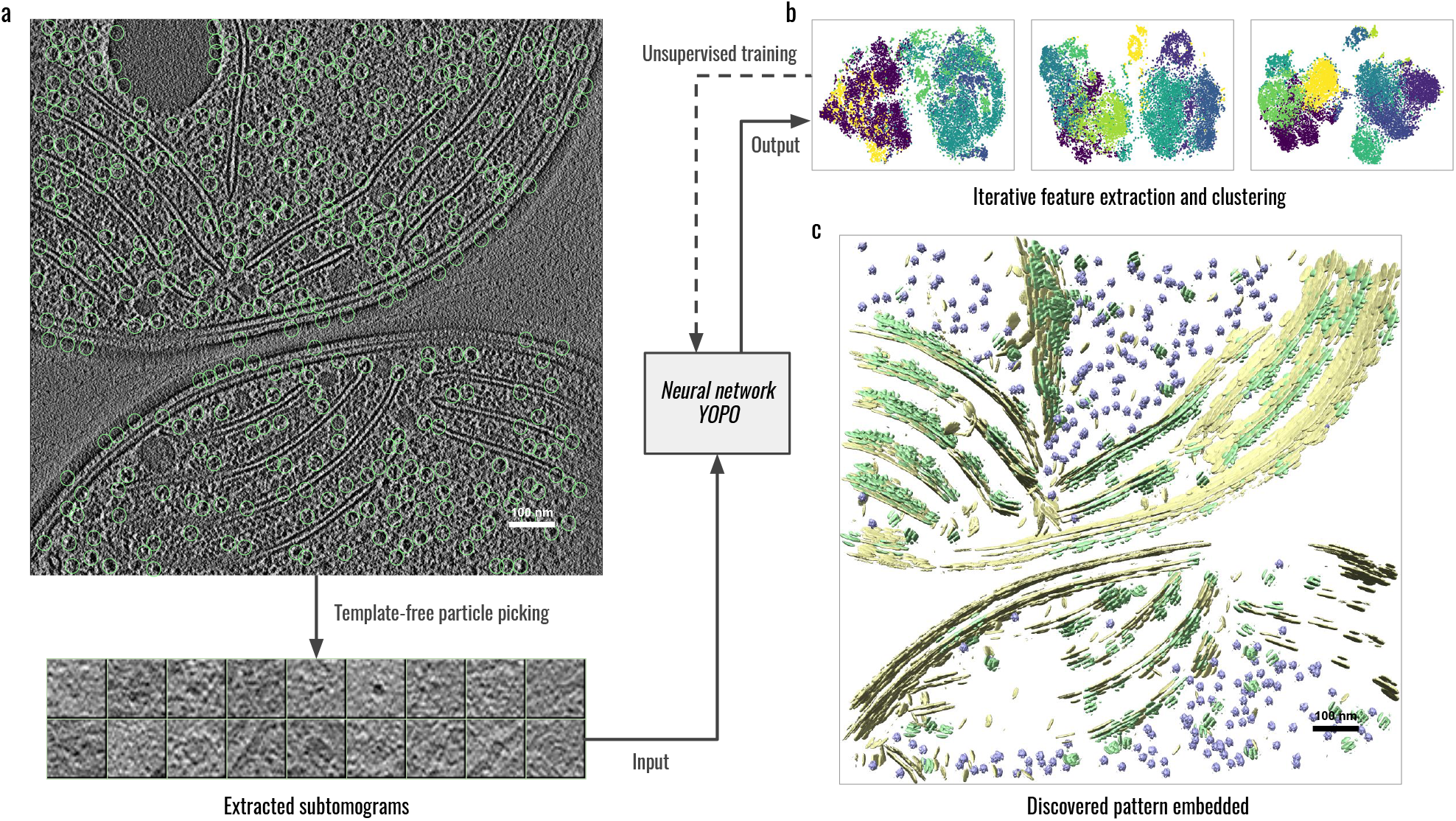
Workflow of DISCA exemplified on a *Synechocystis* cell tomogram (EMDB [13] entry EMD-4603 [14]). (a) 2D slice view of the template-free particle picking on the raw tomogram. (b) Unsupervised training of the YOPO neural network by iteratively clustering extracted features. (c) Discovered patterns by DISCA re-embedded to the original tomogram space.

We tested DISCA on five cryo-ET datasets from distinct cell types (**Fig. 1** and **2**): *Rattus* neuron, *Synechocystis, Cercopithecus aethiops* kidney, *Mycoplasma pneumoniae*, and *Murinae* embryonic fibroblast. Three of the datasets were obtained from public repository EMDB [13] and ETDB [22]. DISCA detected diverse representative structural patterns (detailed results in **Supplementary Note 2**) including macromolecular complexes: ribosome, TRiC, capped proteasome, and phycobilisome array, and other cellular structures: thylakoid membrane, mitochondrial membrane, and calcium phosphate precipitates (**Fig. 2**). The discovered macromolecular complexes have a wide range of sizes from 1.2 MDa to 4.5 MDa in molecular weights. The original manuscripts describing these datasets used manual density selection, template matching, and subtomogram classification to recover the structures. Our unsupervised results from DISCA not only covered the previously identified spatial localization of various macromolecules well but also validated their results in a highly automatic, systematic, and unbiased way. Subtomogram alignment and averaging following DISCA resulted in maps with 14-35 Å resolution range, confirming that template-and-label-free approaches are suited for *in situ* structural analyses.

**Figure 2:**
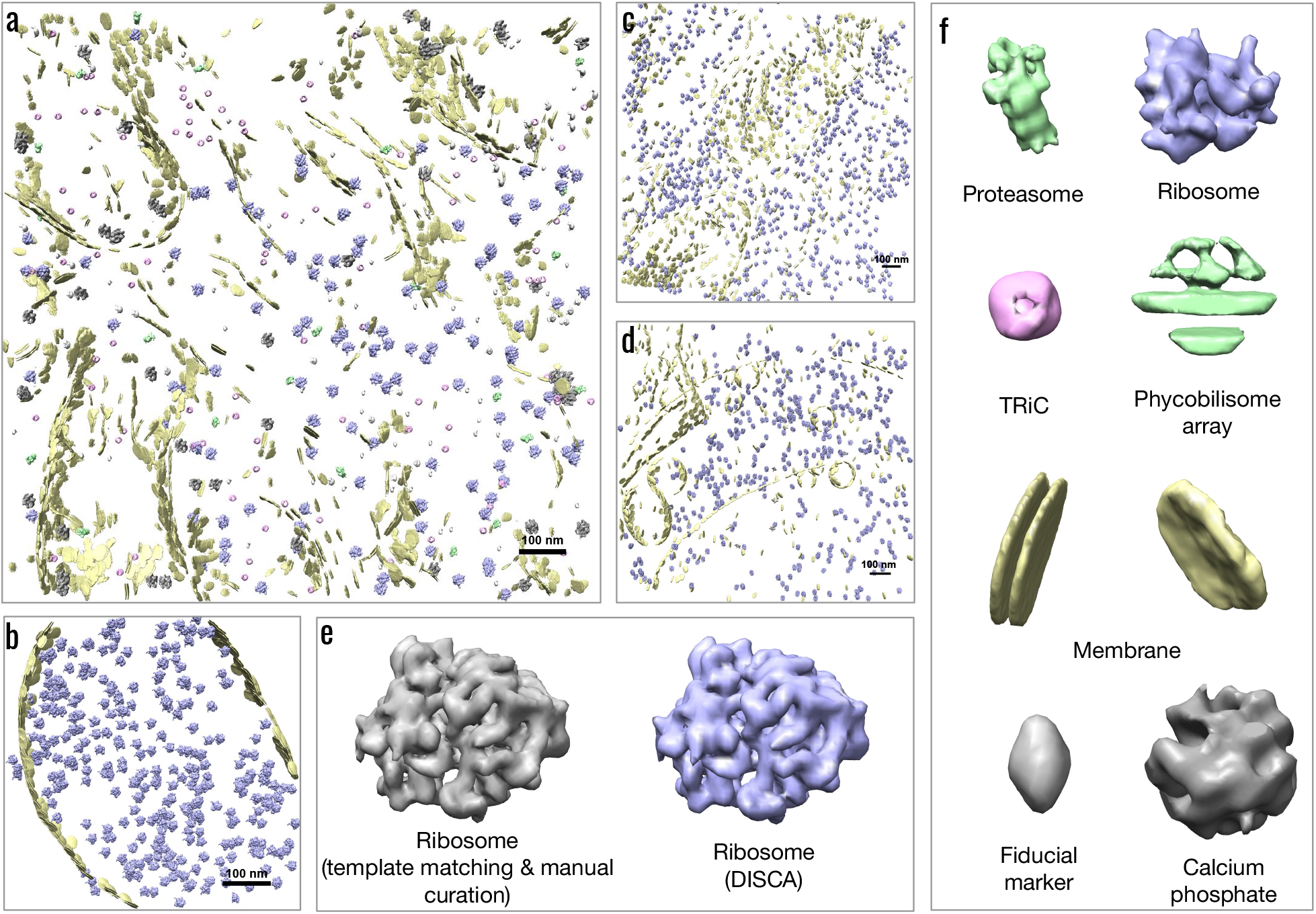
(a-d) Example results of discovered patterns re-embedded to the tomogram of (a) *Rattus* neuron [20]; (b) *Mycoplasma pneumoniae* [21]; (c) *Murinae* embryonic fibroblast [22]; and (d) *Cercopithecus aethiops* kidney [23]. (e) Comparison of recovered ribosome structure from the *Mycoplasma pneumoniae* dataset using template matching (left) and DISCA (right at 14.17 Å). (f) Example results of discovered patterns from the above experimental datasets.

We quantitatively assessed the accuracy of DISCA on the *Mycoplasma pneumoniae* dataset. For this dataset of 65 tomograms, obtaining the clean ribosome particles for comparison required two months of time and heavy computation for traditional 3D template matching, manual curation, and computational sorting. DISCA achieved a low false-positive rate of 9.3% and false-negative rate of 15.0%. Furthermore, DISCA detected about 20% of the ribosomes missed by the template matching and manual curation approach and detected more true ribosomes overall (**Supplementary Note 2**). Notably, DISCA is a very efficient method for processing a large amount of data both theoretically (overall time complexity *O(N)*, where *N* is the number of samples) and practically: on the *Mycoplasma pneumoniae* cell dataset of 65 tomograms, DISCA took less than a day to sort 198,715 template-free picked subtomograms (binned to 24^3^ voxels of 13.33 Å spacing). Moreover, with trained DISCA models, the prediction on new data is very fast and can process millions of such sized subtomograms in less than an hour. The averages of the sorted structural subsets can be further refined to a higher resolution on unbinned data. Even with new data of different image intensities from another source, the trained DISCA models can be fine-tuned without manually preparing training labels. Accordingly, DISCA can efficiently produce meaningful structures from large-scale datasets that encompass very heterogeneous structures without any prior knowledge, which constitutes the first major step for unsupervised structure determination *in situ*. DISCA shows the promise of high-throughput cryo-ET structural pattern mining for discovering abundant and representative structures systematically. The proposed framework will allow researchers to fully leverage state-of-the-art cryo-ET imaging infrastructure and workflows.

## Acknowledgements

This work was supported in part by U.S. NIH grants R01GM134020 and P41GM103712, NSF grants DBI-1949629 and IIS-2007595, and Mark Foundation For Cancer Research 19-044-ASP. J.M. acknowledges support from the EMBL. Y.C. acknowledges support from David and Lucile Packard Fellowship for Science and Engineering (2019-69645). The computational resources were supported by AMD COVID-19 HPC Fund and by Dr. Zachary Freyberg’s lab. X.Z. was supported in part by a fellowship from CMLH. We thank Dr. Qiang Guo and Dr. Tzviya Zeev-Ben-Mordehai for providing testing datasets, Dr. George Tseng for suggestions on the methodology development, and Gregory Howe, Hongyu Zheng, and Dr. Irene De Teresa Trueba for critical comments on the manuscript.

## Author contributions

M.X. and X.Z. conceived the research. X.Z., A.K., and M.X. designed the method. X.Z. implemented and refined the method. L.X. and J.M. provided test dataset. X.Z, L.X., and J.M. evaluated the results. X.Z., Y.C., and M.X. wrote the manuscript. All authors edited the manuscript.

## 2 Methods

DISCA is a generalized Expectation-Maximization (EM) framework that runs iteratively. As shown in **Supplementary Fig. 1** and **2**, in the M-step, the parameters of neural network YOPO are optimized given estimated labels. We describe the architecture design of YOPO and how we achieve rotation and translation invariant feature extraction in the following Methods section. In the E-step, the clustering labels of input subtomograms are estimated using the current parameters based on modeling extracted features. We describe how we statistically model the features and automatically estimate the number of clusters in the following Methods section.

### 2.1 Neural network architecture design

A tomogram is a grayscale 3D volume of very large size (e.g., 4000×6000×1000 voxels). Even binned 4 times across each axis, a tomogram is still large (e.g., 1000×1500×250 voxels). Feeding such a large 3D volume into a CNN will inevitably exceed the memory of the system. One previous CNN method [15] dealt with this problem by slicing the tomogram into 2D images along the z-axis for cost-effective processing. However, taking 2D slices resulted in losing relevant structural information in 3D. In contrast, our objective is to cluster the heterogeneous densities of molecules (the majority being macromolecular complexes) enclosed in subtomograms into structurally homogeneous subsets. Because subtomograms extracted from binned tomograms are significantly smaller (e.g. 24^3^ voxels) than tomograms [24], they can be efficiently processed by 3D CNN without information loss.

Convolutional Neural Networks (CNNs) have been shown to outperform traditional hand-crafted feature extraction methods for the task of extracting discriminative features from images for various biomedical image analysis tasks [25,26]. In order to leverage the superior performance of CNNs, we designed a CNN named YOPO (**Supplementary Fig. 3**) specifically for subtomogram data that considers its distinct characteristics: (1) the structural details are essential to determine the identity of a macromolecule enclosed in a subtomogram; (2) the enclosed macromolecule is of random orientation and displacement; and (3) the Signal-to-Noise Ratio (SNR) is extremely low. Because of the novel architecture design, YOPO achieves properties including structural detail preservation, transformation invariance, and robustness to noise.

#### Structural detail preservation

The standard pooling operation (max-pooling or average pooling) in CNN feature extraction is a problem for processing small 3D subvolumes. Indeed, even pooling by the smallest factor, 2, will dramatically reduce the subvolume size (for example, 24^3^ to 12^3^) and result in losing 87.5% of the information capacity. As structural details predominantly determine a macromolecular complex’s identity, the standard pooling operation is not suitable for extracting features that preserve detailed structural information. Therefore, we equipped YOPO with a sequence of convolutional layers without any pooling operations in between for processing an input subtomogram into feature maps with both low-level and high-level structural information. Following the sequence of convolutional layers, rather than using the basic step of flattening the 3D feature maps into a 1D feature vector, we incorporated a global max-pooling layer to keep only the maximum of each of the feature maps. The global max-pooling operation also achieved translation invariance. As proved later, YOPO will output the same feature values for a subtomogram and its displaced copy because of the translation invariance.

#### Robustness to noise

Another challenge is the extremely low SNR of cryo-ET data. Often, raw tomograms are so noisy that even human eyes barely recognize the structure. While the convolutional layers in YOPO perform filter-like operations, we further boosted YOPO’s robustness to noise. We use a dropout strategy inspired by denoising CNNs to regularize the network and reduce the variance of model prediction from noisy samples. Here, we use a Gaussian dropout layer, which randomly silences 50% of the nodes and injects multiplicative 1-centered Gaussian noise with standard deviation 1 during training. The Gaussian dropout layer has similar regularization performance as the conventional dropout layer, but it exhibits faster convergence properties [27]. By randomly silencing a subset of nodes and injecting Gaussian noise, the Gaussian dropout layer can be viewed as a computationally efficient way to approximate multiple CNNs with slightly different parameters during CNN training. When multiple CNN models are aggregated by inactivating the Gaussian dropout layer during the prediction, the output variance is reduced, thus achieving robustness to noise.

Finally, we added one fully connected layer after the global max-pooling layer to output the feature vectors of length 1024. In order to train YOPO, we equipped the final classification layer with softmax activation to output class labels. The Gaussian dropout layer, self-supervision for rotation invariance, and label smoothing described below have all been shown theoretically and empirically to be effective in preventing overfitting to increase the optimization robustness [28].

### 2.2 Rotation and translation invariant feature extraction

One important characteristic of subtomogram data is that the structure enclosed is randomly oriented and exhibits small random displacement. To cluster multiple copies of the same structure in different orientations and displacements together into the same subset, YOPO must be able to extract features invariant to both translation and rotation.

The rotation invariance was achieved by self-supervised learning for enforcing a CNN to be invariant to certain geometric transformations of the input and improving its generalization ability. In each iteration, alongside the original input subtomogram, a randomly rotated copy of the subtomogram is also fed into YOPO for training. The empty region of the rotated subtomogram is filled with Gaussian white noise. The Gaussian white noise has a mean zero and standard deviation one, same as the normalized image intensity distribution of the input subtomogram data. The label of the randomly rotated copy stays the same. By doing so, the rotation invariance of YOPO is enforced through back-propagating the loss gradient.

The translation invariance is already achieved in the architecture design of YOPO by the global maxpooling layer. The convolution operations *y*_*c*_ are translation equivariant: the extracted feature maps of an input subtomogram *s*_*n*_ translated by *t*_*θ*_ will be the same as translating the extracted feature maps from the original subtomogram by *t*_*θ*_ : *y*_*c*_(*t*_*θ*_ (*s*_*n*_)) = *t*_*θ*_ (*y*_*c*_(*s*_*n*_)). Then, because the global max-pooling layer *y*_*g*_ computes the global maximum from a feature map, which is translation invariant, the output from the global max-pooling layer is translation invariant to the input subtomograms: *y*_*g*_(*t*_*θ*_ (*s*_*n*_)) = *y*_*g*_(*s*_*n*_). Denoting YOPO feature extraction from a subtomogram as: *y*(*s*_*n*_) = *y* _*f*_ ∘ *y*_*g*_ ∘ *y*_*c*_(*s*_*n*_), where *y*_*c*_ denotes the sequence of convolutional layers, *y*_*g*_ the global max-pooling layer, and *y* _*f*_ the fully connected layer, we have:

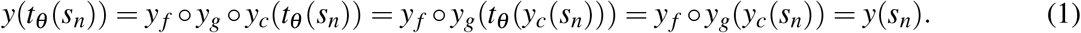

As a result, the final extracted feature vectors are translation invariant to the input subtomograms. We also empirically verified the transformation invariance learned by YOPO (**Supplementary Note 1**).

When designing YOPO, we have tested alternative architectures such as 3D InceptionNet and ResNet as feature extractors, and incorporated other layers including max-pooling, average pooling, global average pooling, flatten, and conventional dropout layers into the network design. The final YOPO design was based on empirically comparing alternative architectures.

### 2.3 Statistical modeling of structurally homogeneous subsets in feature space

Recent works [29,30] have shown that second-order statistics in CNNs—for instance, the covariance between features—are vital for differentiating between different visual patterns. Accordingly, simple clustering algorithms such as K-means or hierarchical clustering which do not consider second-order statistics are not suitable. To fully capture the feature covariance information, after extracting the translation and rotation invariant features from the input subtomograms by YOPO, we model the learned feature vectors for each representative structural pattern as a multivariate Gaussian distribution in the feature space.

In greater detail, given a set of *N* subtomograms *s*_*n*_ ∈ *S* extracted from a dataset of tomograms *V*, the YOPO network *y* extracts feature vectors *x*_*n*_ = *y*(*s*_*n*_), *x*_*n*_ ∈ ℝ^*P*^ from each subtomogram, where *P* is the dimensionality of the feature space. We model the distribution of the data point *x*_*n*_ as a mixture of *K* multivariate Gaussian distributions. The mixture distribution’s probability density *f*_*g*_ is defined as:

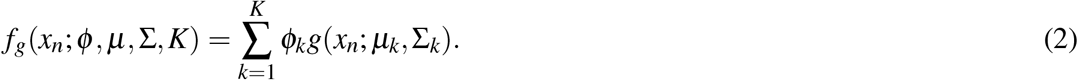

In Eq. 2, *ϕ*_*k*_ is the prior probability of sampling *x*_*n*_ from the *k*th component. The *k*th component is a multivariate Gaussian distribution *g* with mean *µ*_*k*_ and covariance matrix Σ_*k*_. Hence, the posterior probability of sampling *x*_*n*_ from the *k*^*th*^ component is 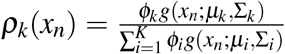. Solving the model [31] in Eq. 2 provides the probability *ρ*_*k*_(*x*_*n*_) of feature vector *x*_*n*_ being sampled from each component distribution *g*(*x*_*n*_; *µ*_*k*_, Σ_*k*_). *g*(*x*_*n*_; *µ*_*k*_, Σ_*k*_) has its own covariance matrix Σ_*k*_ to distinguish between different structural patterns. The component 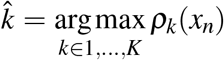 is the highest posterior probability among all components. 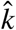 will be used as the class label for subtomogram *s*_*n*_ in the clustering solution.

### 2.4 Automatic estimation of the number of structurally homogeneous subsets

Because we operate in an unsupervised learning setting, the number of structurally homogeneous subsets *K* is unknown to us. Furthermore, the automatic estimation of the number of clusters in a feature space is a classic yet highly challenging and largely unsolved problem, which means that, practically, most studies just set an arbitrary *K* or test multiple candidate values of *K* and manually compare the results. Nevertheless, in our statistical modeling, it is beneficial to choose *K* properly. When the chosen *K* is too small, a subset may contain mixed structures. In contrast, when the chosen *K* is too large, a structurally homogeneous subset may be over-partitioned to multiple subsets. Over-partitioning likely results in some subsets containing too few subtomograms to recover the structure. Both situations may lead to poorly recovered structures by subtomogram averaging. For this reason, it is helpful to automatically determine *K*.

Automatic estimation of *K* relies on observing the extracted feature vectors. Most recent and popular methods for estimating *K* are either prediction-based or stability-based, and require running the given clustering algorithm repeatedly on bootstrapped samples [32]. These methods are not suitable for our study because they are too slow to process large-scale datasets. Other methods for estimating *K* compute a summary index measuring cluster tightness. For example, the silhouette coefficient compares the average distance of a data point to all the other data points in its own cluster and in its nearest cluster. However, computing the silhouette coefficient involves comparing all pairs of data points (time complexity: *O*(*N*^2^)), which is still poor in scalability.

To overcome these shortcomings, we take an alternative approach from a statistical model selection perspective. The number of model parameters increases along with *K*, which may result in increased likelihood, but also runs the risk of overfitting. When modeling the structurally homogeneous subsets in the feature space, a good statistical model would ideally have a higher likelihood with relatively few parameters. To balance the likelihood and number of parameters among a set of models with different *K*s, we use the Bayesian Information Criterion (BIC) [33] to select among a set of fitted models *M*, where the BIC is defined as:

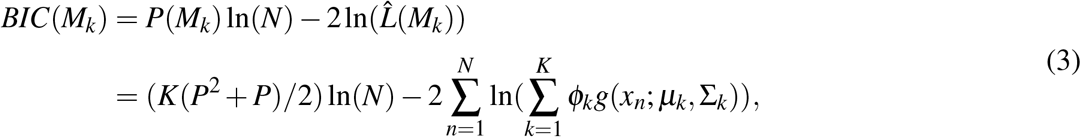

where *M*_*k*_ is the fitted model with *K* structurally homogeneous subsets, *P*(*M*_*k*_) denotes the number of parameters in model *M*_*k*_ and 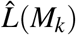 denotes the maximized value of the likelihood function of *M*_*k*_. The model with the lowest BIC is selected. To validate the model selection, we conducted several experiments on simulated datasets of various SNR and tilt-angle ranges and included experimental comparison with baseline methods (**Supplementary Note 1**). We also tested Akaike information criterion (AIC) [34], CH index [35], KL index [36], and Jump statistic [37], our preliminary results showed that BIC achieved superior performance.

### 2.5 Iterative dynamic labeling

In supervised learning, a CNN is trained to maximize the prediction accuracy on a set of labeled training data. As we only have unlabeled data, we develop a strategy to iteratively estimate both the number of structurally homogeneous subsets and the structural class labels of input subtomograms. The proposed iterative dynamic labeling strategy updates two models in an alternating fashion via a generalized Expectation-Maximization (EM) algorithm [38]. **Supplementary Fig. 3** illustrates the YOPO model for feature extraction and the Gaussian distributions for the statistical modeling of structurally homogeneous subsets in the feature space ℝ^*P*^. In the E-step, the number of structurally homogeneous subsets and the labels are estimated given the current learned features according to Eq. 2. In the M-step, YOPO parameters are updated by back-propagation training to minimize the loss function of computing the labels estimated from the E-step. For the optimization of the statistical model fitting, it is stabilized by inheriting the parameters from the previous iteration. Moreover, because errors can accumulate when initializing the statistical model fitting using parameters from the previous iteration, to avoid getting stuck at a local optimum, a *de novo* model fitting with randomly initialized parameters was also performed in each iteration and its parameters were adopted if this model increased the likelihood.

In summary, YOPO is randomly initialized to extract feature vectors *x*_*n*_ ∈ ℝ^*P*^ from input subtomograms *s*_*n*_ ∈ *S*. Then, the feature vectors are fitted in the feature space by the mixed multivariate Gaussian distributions across a set of candidate *K* number of structurally homogeneous subsets. Only the mixture distribution with the lowest BIC is kept. The current estimated label of a subtomogram is given by a hard cluster assignment that corresponds to the component multivariate Gaussian distribution with the highest probability. In the next iteration, the current estimated labels are used for training YOPO by minimizing the categorical hinge loss function to learn better feature representations. After YOPO training, the mixture distributions is updated on the newly extracted feature vectors by optimizing Eq. 2. This process continues iteratively until convergence.

A potential issue is that, unlike in supervised learning, where training data labels are fixed, the YOPO training data labels are dynamic. In other words, there will inevitably be mislabeled data when training YOPO, especially in the early iterations. To address this issue, we adapt the label smoothing regularization technique [39] to make the YOPO training less prone to mislabeled data. The smoothed one-hot encoding of training labels is: 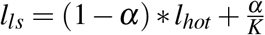, where *K* is the number of clusters, *l*_*hot*_ is the original one-hot encoding of training labels, and *α* is the smoothing factor. The larger the label smoothing factor *α*, the less certain the model prediction.

Moreover, the estimated *K* is also dynamic in different iterations. We need to enable YOPO to output different class numbers during the training in different iterations. When the estimated *K* differs from the previous iteration, we replace the last layer, the classification layer, with a new one with the current estimated *K* number of nodes. Because the new classification layer has randomized initial weights, we train its weights with the fixed current extracted features as input to reach consistency between its prediction and current estimated labels.

### 2.6 Matching clustering solutions

From our experience (**Supplementary Note 1**), the estimated *K* stays the same in most iterations. In such cases, instead of replacing the last classification layer, we directly match the current clustering solution with the one in the previous iteration. When there are multiple clustering solutions from the same samples, the label of a specific cluster is not necessarily the same between different solutions. For example, the same group of samples may be labeled as ‘1’ by one clustering solution and ‘2’ by another even if they result from the same clustering algorithm with exactly the same parameters. The inconsistency will cause strong instability during training (**Supplementary Fig. 13**). Therefore, matching clustering solutions is necessary.

We formulate the problem of matching two clustering solutions as a maximum weighted bipartite matching problem. First, we define a bipartite graph that consists of two disjoint and independent sets. In our case, the two sets are the two clustering solutions from consecutive iterations. Then, we define a cluster as a graph vertex and the number of overlapping samples in two vertices (one in each of the two clustering partitions) as the graph edge weight. Maximum weighted bipartite matching finds a subset of the edges where no two edges share a common vertex and maximizes the sum of edge weights. In our case, the two sets have the same number of vertices (*K*) and each vertex has precisely one edge in the optimal matching.

Let *B* be a Boolean matrix to represent the matching where *B*_*i, j*_ = 1 if cluster *i* in *a* is matched to cluster *j* in *b*. The optimal matching is formulated by maximizing the objective function:

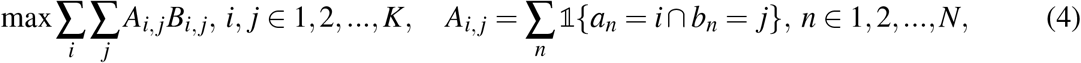

where *A* is the matching matrix (a.k.a. confusion matrix in supervised learning) between the two solutions *a* and *b*, and 𝟙{} is the indicator function.

In each iteration of DISCA, the estimated labels are assigned on model fitting solutions to Eq. 2. Due to the reasons mentioned above, in DISCA, the clustering solution from one iteration needs to be matched with the previous clustering solution to stabilize the training. We apply the Hungarian algorithm [40] to optimize the objective function (Eq. 4), which is guaranteed to find a global optimum in polynomial time. Then, the current labels are permuted according to the matching to achieve the highest consistency with the labels in the previous iteration.

### 2.7 Missing wedge effect

A major cryo-ET limitation, the missing wedge effect, must be considered when designing analysis methods [41]. In cryo-ET imaging, cell samples are imaged through a series of tilt projections. The tilt projections are subsequently fed into a reconstruction algorithm to produce a 3D tomographic reconstruction. Because of the increasing effective sample thickness during tilting, to prevent excessive electron beam damage to the cell sample, the tilt angle range is limited typically to ±60° with a 1° step size. This results in a double V-shaped missing value region of Fourier coefficients of the reconstructed tomogram in Fourier space. The missing wedge effect also produces image distortion in the spatial domain; for instance, it may elongate features along the direction of the missing wedge axis.

DISCA tackles the missing wedge effect from two aspects. First, in our previous work [23], we have empirically demonstrated the robustness of CNN feature extraction to image distortions caused by the missing wedge effect. Moreover, the robustness of YOPO feature extraction to image noise and distortion is further improved by the Gaussian dropout layer. Second and most importantly, during the self-supervision step, when a subtomogram is rotated, the direction of the image distortion caused by the missing wedge effect rotates correspondingly. By enforcing the rotated copy to have the same label and thus similar extracted feature vectors during YOPO training, we explicitly increase the robustness of YOPO feature extraction to the missing wedge effect from various angles. In **Supplementary Note 1**, we showed that DISCA can still perform well on simulated datasets of large missing wedge (tilt-angle range ±40°) and various SNR, thus demonstrating the robustness of DISCA to the missing wedge effect.

The missing wedge effect can also be treated in other data processing steps. Before feeding into DISCA, the tomograms can be reconstructed by algorithms compensating for the missing wedge effect such as Weighted BackProjection. In the postprocessing step, subtomogram averaging using *RELION* [42] involves missing wedge compensation from model estimation, whereas structural pattern re-embedding by Gum-Net [17] uses a spectral data imputation technique to reduce the missing wedge effect on subtomogram alignment.

### 2.8 Time cost and complexity analysis

Currently, there are more than 100 TB of cryo-ET data in public repositories such as EMDB [13], ETDB [22], and EMPIAR [43]. With the fast accumulation of cryo-ET data, it is necessary to have high-throughput analysis algorithms. We now show theoretically that DISCA can achieve an overall time complexity of *O*(*N*), and therefore our framework scales well to large datasets. This leads to the following theorem.

#### Theorem 1.

*When m, the number of iterations, K, the number of clusters, and P, the dimension of the feature space, are held constant and are relatively small compared to N, the number of entries in the dataset, the time complexity of DISCA is O*(*N*).

*Proof*. In each of *m* iterations, the algorithm performs feature extraction by YOPO, estimates the number of components, fits mixed multivariate Gaussian distributions to the extracted features, matches clustering solutions, validates clustering solutions, and trains the YOPO network using current estimated labels. The deep learning process to extract features takes time *O*(*N*). Estimating *K* using BIC takes time *O*(*K*). Statistical model fitting takes time *O*(*NKP*^2^) using the FIGMN algorithm [44]. In the matching stage, the Hungarian algorithm takes time *O*(*K*^3^) [40]. Finally, when validating clustering solutions, calculating the distortion-based DBI takes time *O*(*N*).

Therefore, the total time complexity is *O*(*m*(*N* + *K* + *NKP*^2^ + *K*^3^ + *N*)), but because *m, K*, and *P* are constant, the overall computational complexity of DISCA is *O*(*N*).□

In terms of sample complexity, we leverage work by [45] that has shown that 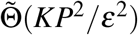 samples are both necessary and sufficient for learning mixed multivariate Gaussian distributions with *K* components in a *P*-dimensional feature space with up to *ε* error in total variation distance. This result implies that learning reasonably accurate models that achieve a low, constant error *ε* requires relatively few samples in practice, as *K* and *P* are assumed to be small compared to *N* in large-scale datasets.

Practically, on our computer with 4 GPUs and 48 CPU cores, the pre-processing template-free particle picking step takes less than 20 minutes to pick 100,000 to 200,000 subtomograms from a dataset of more than 10 tomograms. Training DISCA from scratch to sort these subtomograms takes less than 10 hours. When our clustering model is properly trained, the prediction on new data is very fast, which takes less than an hour to process millions of subtomograms. Before the subtomogram averaging step, the cluster centers of extracted features can optionally be decoded to select interesting clusters for thorough downstream analysis (**Supplementary Note 3**). The post-processing subtomogram averaging step using *RELION* [42] takes less than two days to achieve resolution better than 35 Å. Here, we use ‘subtomogram averaging’ to refer to the averaging process to recover a single class and ‘subtomogram classification’ to refer to averaging and classification process to recover multiple classes which is more time-consuming. By comparison, the template matching approach on the same computer equipment would take roughly one to two months to complete, which requires visual inspection by experts, computational template matching, and subtomogram classification.

## Data source

The *Rattus* neuron dataset is obtained from [20]. The *Synechocystis* dataset is obtained from EMDB [13] entry EMD-4603 and EMD-4604 [14]. The *Cercopithecus aethiops* kidney dataset is obtained from [23]. The *Murinae* embryonic fibroblast is obtained from ETDB [22] with MefB cell line from O. Loson in Chan Lab. The *Mycoplasma pneumoniae* dataset was acquired as described previously [21]. Tomograms were reconstructed and filtered using *Warp* [46]. The original tilt-series data is available via EMPIAR-10499.

## Supplementary Information

**Figure S1:**
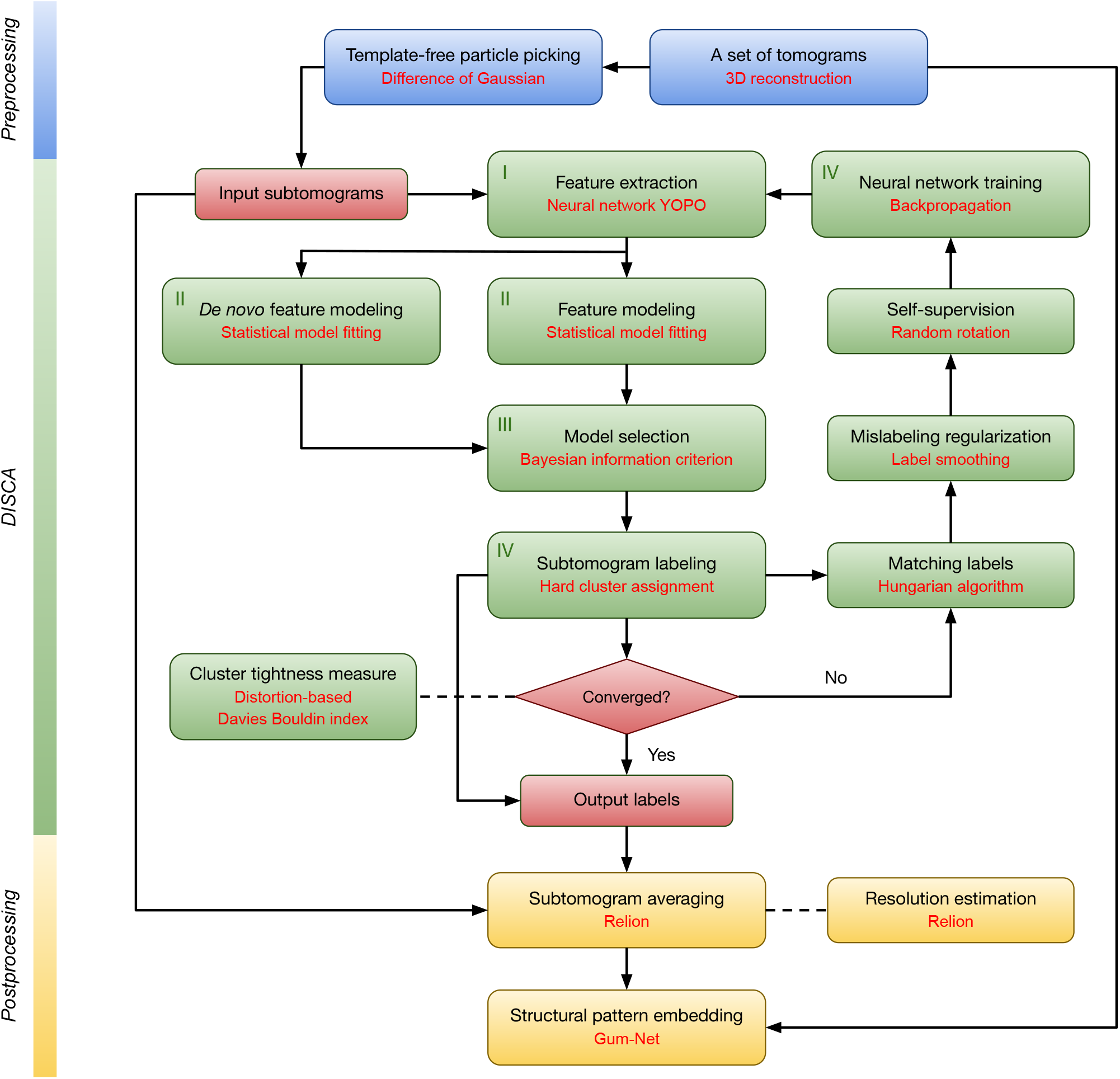
The DISCA workflow for cryo-ET structural pattern mining. Key steps are numbered. The preprocessing and postprocessing steps are included here for an overview of the processing pipeline. They are not part of the proposed method DISCA.

**Figure S2:**
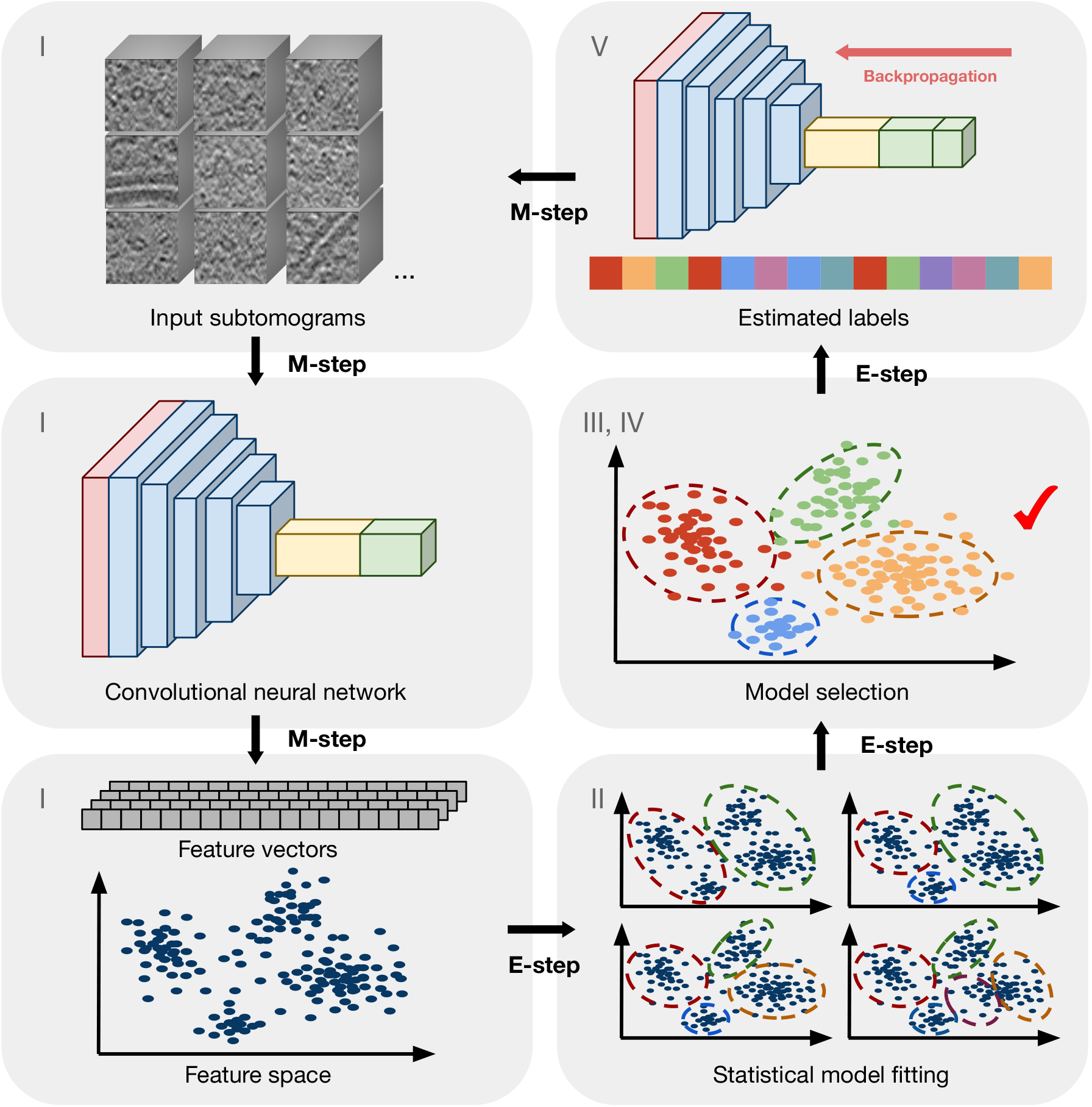
Conceptual explanation of DISCA. The numbers correspond to key steps in **Fig. S1**. The input is a set of subtomograms extracted from tomograms using template-free picking methods. CNN features extracted (step I) from subtomograms are statistically modeled (step II) to estimate the cluster labels (step II and IV). The CNN is in turn trained (step V) using the current estimated labels in order to learn better features iteratively.

**Figure S3:**
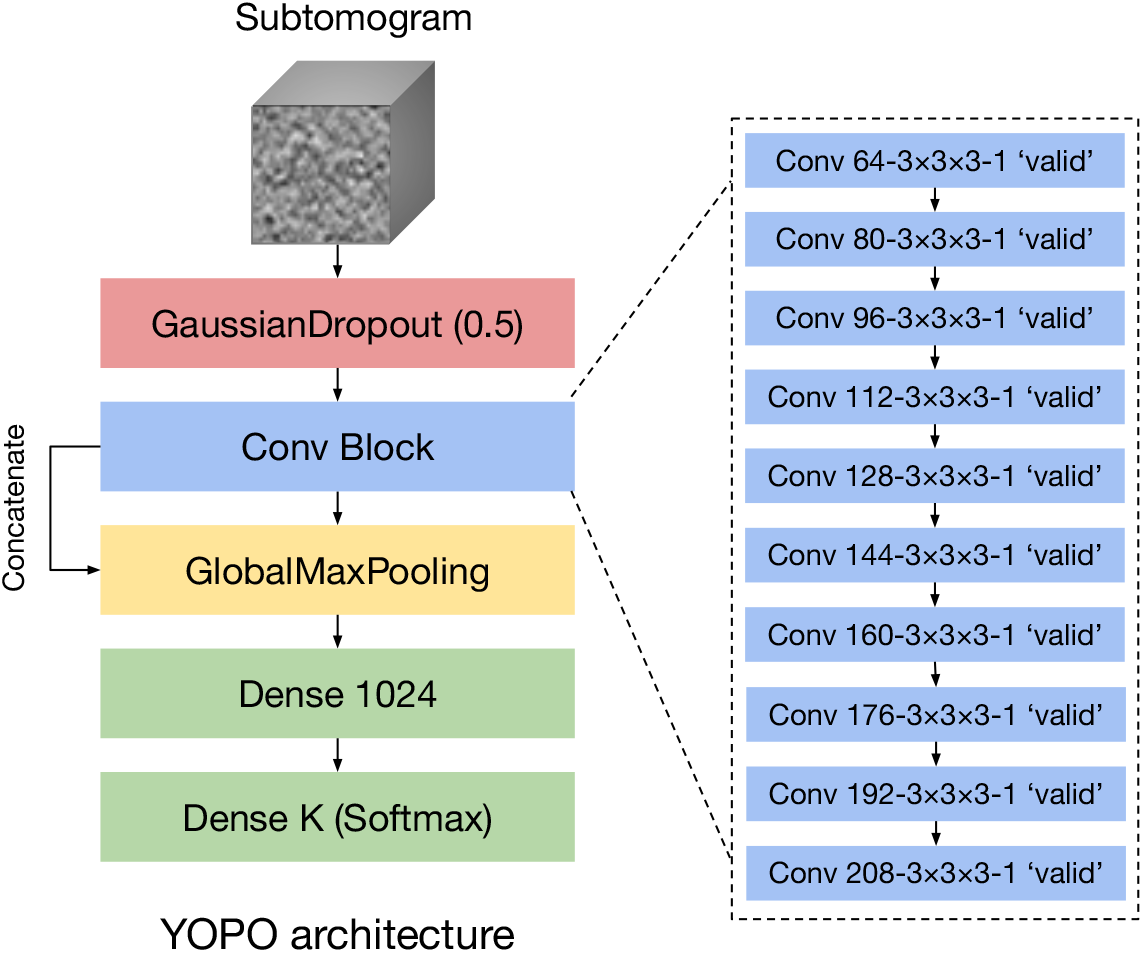
The architecture of YOPO (You Only Pool Once) model. Each colored box denotes one layer in the neural network. ‘GaussianDropout (0.5)’ denotes a dropout layer with dropout rate of 0.5 and multiplicative 1-centered Gaussian noise. ‘Conv 64-3×3×3-1 ‘valid” denotes a convolutional layer with 64 channels, kernel size 3 × 3 × 3, strides of size 1, and valid padding (no padding). Each convolutional layer is equipped with exponential linear unit activation function and batch normalization. ‘Concatenate’ denotes concatenated feature outputs. ‘Dense K (Softmax)’ denotes a fully connected layer with K neurons. The extracted features are the output from the ‘Dense 1024’ layer.

### Supplementary note 1: validation of the feature learning and modeling ability of DISCA

To validate DISCA’s ability to learn to extract and model 3D transformation-invariant features, we conducted several experiments on realistically simulated datasets, which have pre-specified ground truth labels to quantitively assess the performance of DISCA and existing methods.

To test the accuracy of DISCA in simultaneously estimating the number of clusters *K* and structural class labels, we simulated subtomogram datasets of various SNR and tilt-angle ranges. We used a standard subtomogram simulation procedure [1,2] and took into account the tomographic reconstruction process with missing wedges and a contrast transfer function. The simulated imaging condition is similar to real experimental settings [3] with voltage 300 KeV, defocus −5 *µ*m, and spherical aberration 2.7 mm. We chose five representative macromolecular structures: RNA polymerase (1I6V), rotary motor in ATP synthase (1QO1), proteasome (PDB ID: 3DY4), ribosome (4V4A), spliceosome (5LQW). Real cryo-ET data typically have an SNR below 0.1 [4] and a tilt-angle range around −60° to 60°. For each macromolecular structure, we simulated 400 subtomograms at each SNR (0.1, 0.03, 0.01, 0.003, and 0.001) and tilt-angle range (±60° and ±40°) to demonstrate the robustness of DISCA to the image noise and the missing wedge effect.

#### Estimating *K* and labels

We performed DISCA on each of the simulated dataset. We evaluated the results by three criteria: (1) the estimated *K* with candidate *K* ranging from 2 to 20; (2) the homogeneity score measuring how homogeneous each cluster is according to the ground truth labels. We note that the homogeneity score does not require equal number of clusters to the ground truth; (3) the prediction accuracy measuring the percentage of correctly labeled subtomograms. The prediction accuracy can only be calculated when *K* is estimated correctly. The results from **Table S1** show that DISCA correctly estimated the true *K* for eight of the ten datasets except at SNR 0.003 and 0.001 of tilt-angle range ±40°. As expected, the homogeneity scores gradually decreased with lower SNR and smaller tilt-angle range. However, in all settings, we achieved good results with homogeneity scores higher than 0.8, which means that the resulting clusters are generally homogeneous. We have conducted the experiments using randomly initialized models multiple times. The results were similar with ±5% margin, which ensured the reproduciblity of our method.

We additionally performed template matching and autoencoder clustering for comparison. For template matching, even though we incorporated prior domain knowledge of known structural templates and thus *K*, the results are still worse than DISCA because template matching is not robust to noise. Under SNR lower than 0.01, template matching failed with accuracy close to random guess (20%). We previously proposed the first unsupervised deep learning model to cryo-ET data [5], a convolutional autoencoder that coarsely groups and filters raw subtomograms. In that paper, we proposed a pose normalization step to normalize the orientation and displacement of structure inside a subtomogram for better structural grouping. Compared with DISCA, the convolutional autoencoder can only perform coarse grouping with homogeneity score lower than 0.55. This is mainly because DISCA is a significantly more sophisticated method which involves iterative feature learning and modeling in order to recognize the fine structure differences between different types of macromolecules.

**Table S1:** Performance of three methods on simulated datasets. In each cell, the first row denotes the estimated *K* for unsupervised methods. The second row denotes homogeneity score compared to ground truth. The third row denotes prediction accuracy.

| Dataset | Simulated $\pm 60^\circ$ | | | | | Simulated $\pm 40^\circ$ | | | | |
| --- | --- | --- | --- | --- | --- | --- | --- | --- | --- | --- |
|  | SNR 0.1 | 0.03 | 0.01 | 0.003 | 0.001 | SNR 0.1 | 0.03 | 0.01 | 0.003 | 0.001 |
| Template Matching | - | - | - | - | - | - | - | - | - | - |
|  | 0.7013 | 0.4709 | 0.1496 | 0.0136 | 0.0032 | 0.5543 | 0.3336 | 0.0655 | 0.0062 | 0.0012 |
|  | 83.95 % | 69.75 % | 45.35 % | 25.25 % | 20.95 % | 76.25 % | 61.15 % | 36.60 % | **23.80 %** | **21.20 %** |
| Autoencoder | K = 5 | K = 4 | K = 5 | K = 5 | K = 3 | K = 6 | K = 5 | K = 3 | K = 3 | K = 3 |
|  | 0.3843 | 0.4539 | 0.3613 | 0.4915 | 0.3881 | 0.5227 | 0.3470 | 0.3735 | 0.3878 | 0.3874 |
|  | 56.75 % | - | 53.45 % | 64.80 % | - | - | 53.35 % | - | - | - |
| DISCA | K = 5 | K = 5 | K = 5 | K = 5 | K = 5 | K = 5 | K = 5 | K = 5 | K = 6 | K = 6 |
|  | **0.9878** | **0.9373** | **0.8746** | **0.8712** | **0.8719** | **0.9568** | **0.8020** | **0.8344** | **0.8366** | **0.8323** |
|  | **99.70 %** | **97.80 %** | **94.80 %** | **94.25 %** | **94.50 %** | **98.70 %** | **90.35 %** | **91.80 %** | - | - |

In addition, we provided a summary index, modified from the Davies-Bouldin Index (DBI) [6], as an indicator measuring the cluster tightness relative to cluster separation. Rather than using Euclidean distance in the feature space, we used a distorted measure of the distance which takes each cluster’s covariance into account. We mathematically formulate the proposed distortion-based DBI (DDBI) as:

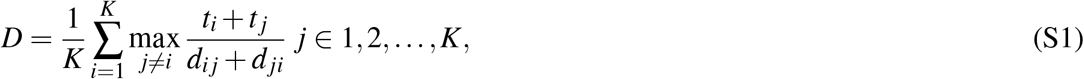

where *t*_*i*_ measures the tightness of *i*th cluster (same for *t* _*j*_) and *d*_*i j*_ measures the separation between cluster *i* and *j*:

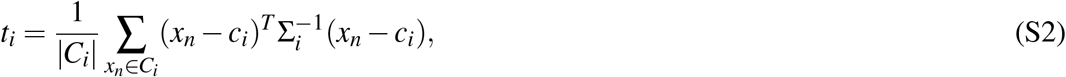

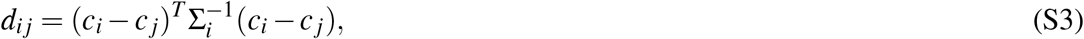

where *C*_*i*_ denotes the subtomograms *x*_*n*_ in the *i*th cluster and *c*_*i*_ denotes its centroid.

We further conducted several experiments and demonstrations using simulated dataset SNR 0.01 and tilt-angle range ±60°, which is closest to the image condition of real datasets as measured on the *Synechocystis* cell [7] and *Rattus* neuron [3] tomograms. In **Fig. S4**, *K* was estimated at 4 for early iterations, where some clusters were not separated well. Extracted features gradually separated out through the iterative learning process. We achieved lowest DDBI at iteration 15, which was kept as the final result.

**Figure S4:**
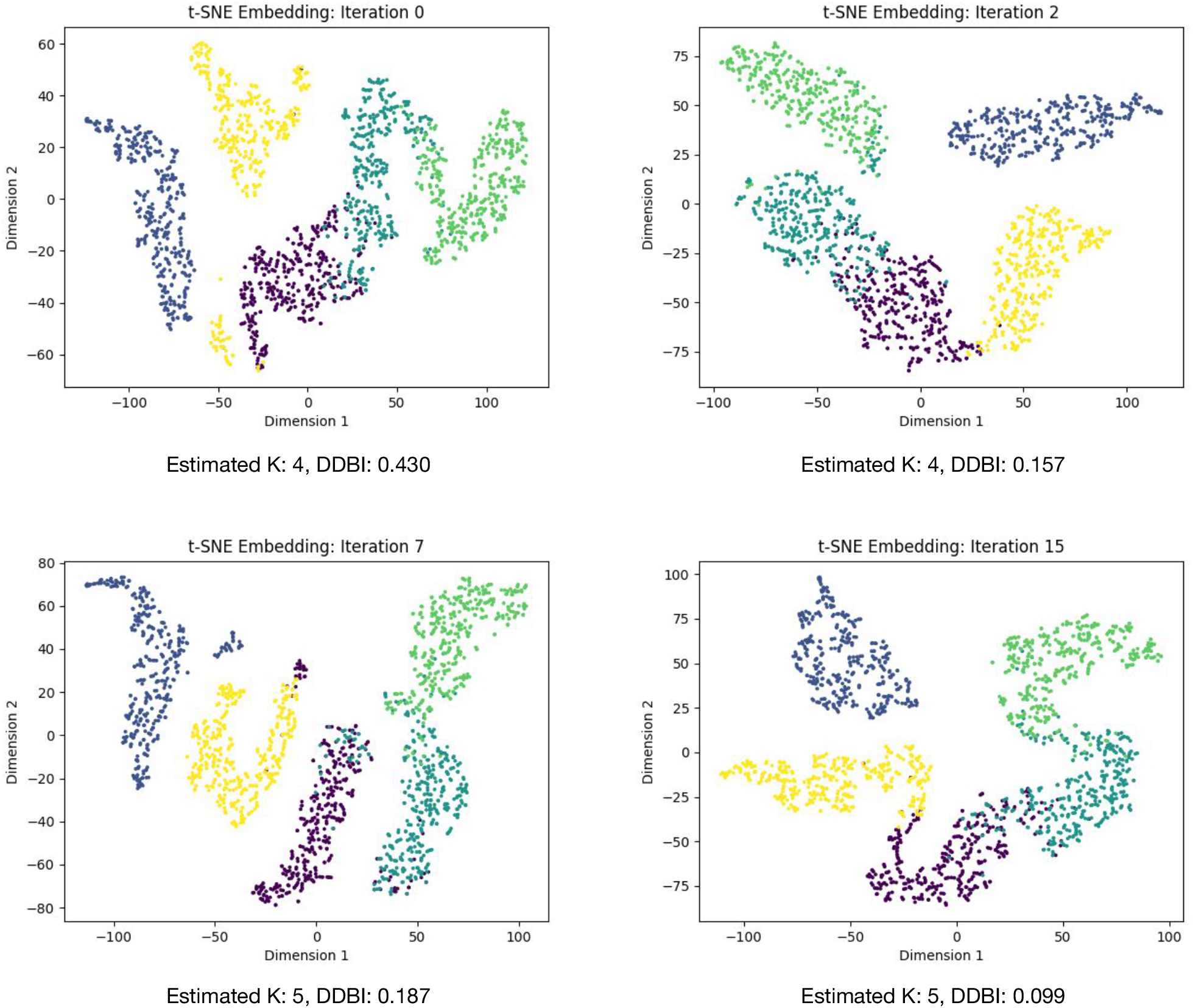
T-SNE [8] embedding of extracted features in different iterations. Each dot denotes one sample with its color indicating its structural class.

#### Progressively improved performance with larger sample size

To demonstrate the learning ability of DISCA with respect to different sample size, we conducted experiments varying input subtomogram number from 50 (10 subtomograms of each structural class) to 10,000 (2,000 subtomograms of each structural class). The results are shown **Fig. S5**.

**Figure S5:**
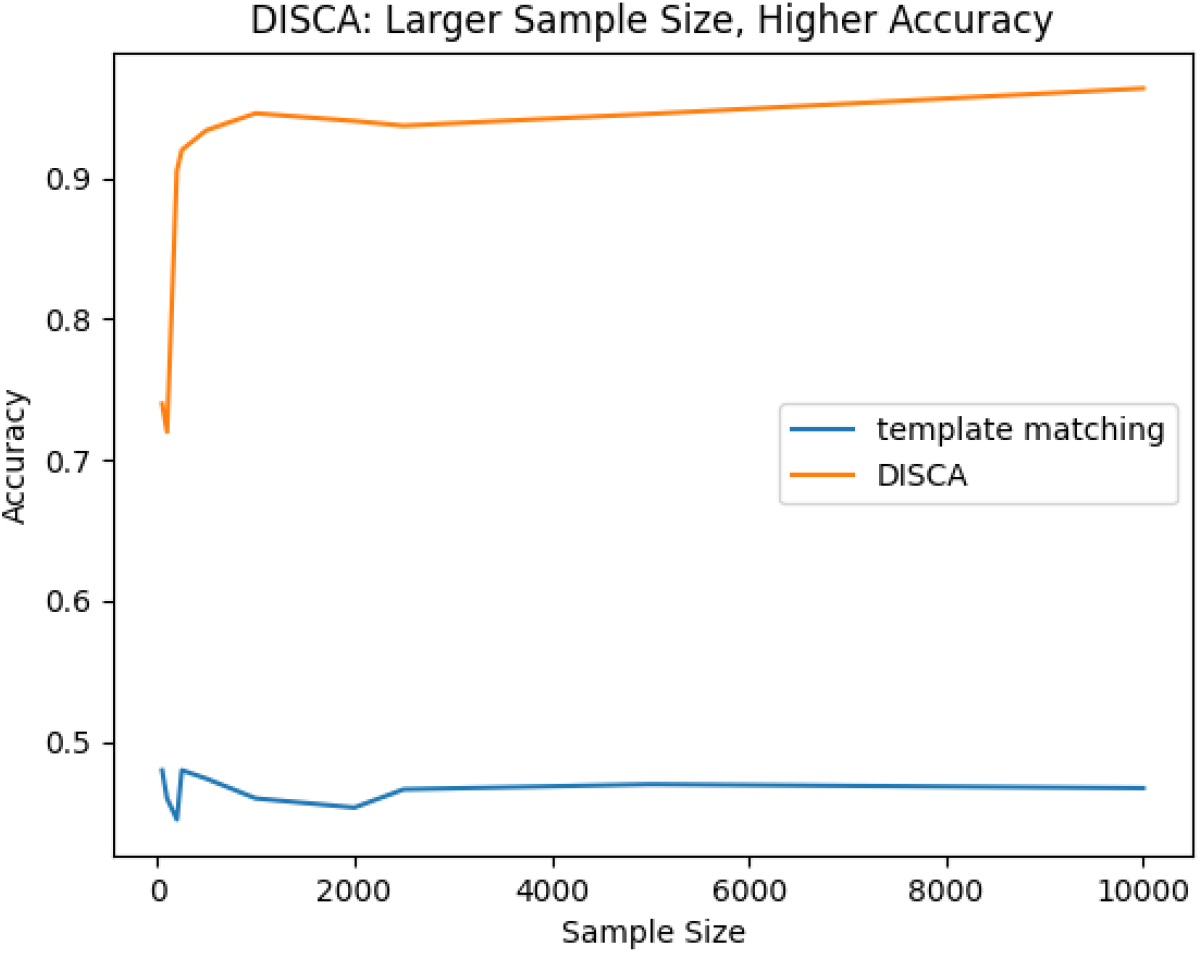
Accuracy of template matching and DISCA with respect to different sample sizes. The accuracy of DISCA improves progressively with larger sample size. The accuracy of template matching stays the same because it does not involve a learning process.

#### Transformation-invariant feature extraction

To verify that the trained YOPO model extracts 3D features that are transformation-invariant to a large extent, we randomly chose one subtomogram from each class and generated 1,000 randomly rotated and translated copies. The extracted features are visualized in **Fig. S6**. We can see that features extracted from transformed copies are very similar to each other as compared to transformed copies of subtomograms of other classes.

**Figure S6:**
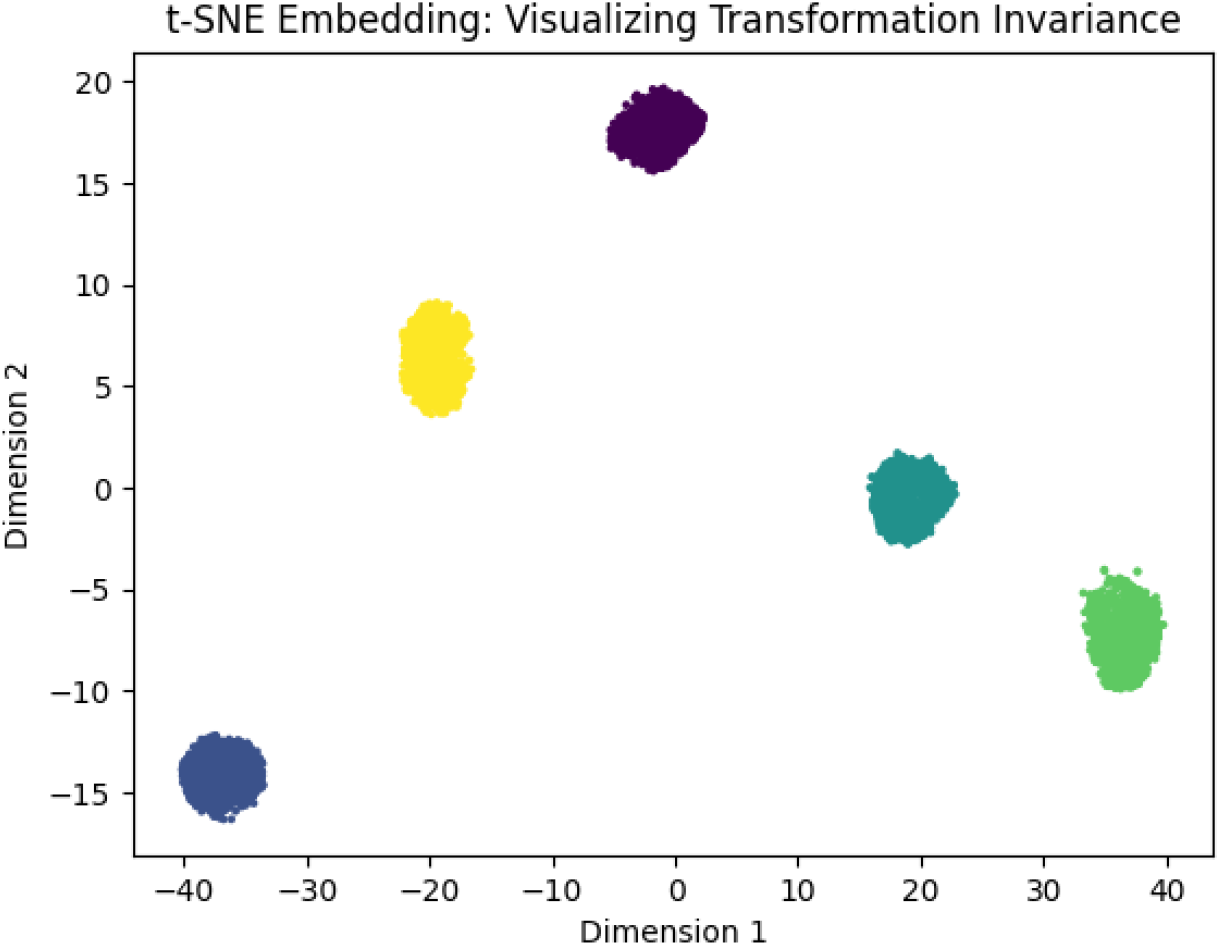
t-SNE embedding of extracted features from randomly transformed subtomogram copies (1,000 copies per subtomogram). Each dot denotes one copy with its color indicating its structural class.

### Supplementary Note 2: detailed results

**Fig. S7** compares example raw tomogram slices and the corresponding re-embedding annotations of discovered patterns from a set of 65 *Mycoplasma pneumoniae* cellular tomograms [9]. The voxel size of this tomogram is 6.802 Å and the resolution measured on the ribosome pattern is 14.17 Å. For comparison, we applied template matching, manual curation, subtomogram classification by *RELION* [10] to recover the ribosome structure, which is referred to hereafter as the *template matching approach*. We consider two detections as overlapping if their Euclidean distance is smaller than 8 nm. Under this criterion, 96.9% of the 18,987 ribosomes detected by template matching are included in the 198,715 subtomograms extracted by template-free particle picking.

**Figure S7:**
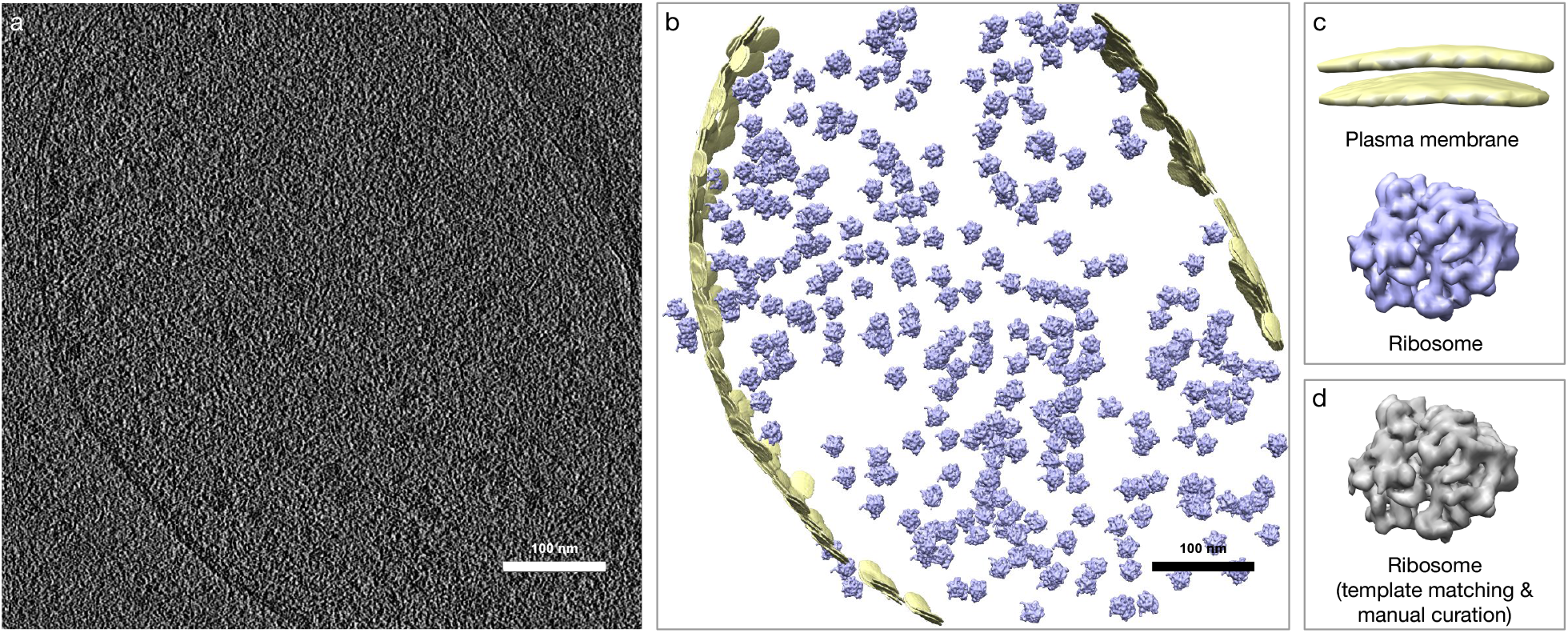
Example unsupervised annotation on a *Mycoplasma pneumoniae* cell tomogram [9]: a. slice of the original tomogram. b. discovered patterns re-embedded to the original tomogram space. c. iso-surface visualization of discovered patterns identified (generated from subtomogram averaging). d. iso-surface visualization of the ribosome structure using the template matching approach.

DISCA clustered the 198,715 total extracted subtomograms into ten clusters where one cluster corresponds to ribosome structures and one cluster corresponds to membrane structures. Among those ribosomes detected by template matching, 85.0% of them overlap the ribosome clusters and consist of 22,875 subtomograms. On the other hand, 70.4% of the 22,875 ribosomes detected by DISCA overlap with the template matching results. As shown in **Fig. S7** c and d, the template-and-label-free result from DISCA resembles the template matching result with correlation coefficient 0.995.

We further investigate the 6,768 ribosomes uniquely detected by DISCA. To assess how many of them are truly ribosomes, we used *RELION* subtomogram classification function to classify them into 4 classes. We note that we did not apply the traditional template matching method to identify them because these ribosomes were missed by the template matching approach. As shown in **Fig. S8**, class 1, 2, and 3 clearly correspond to the ribosome structure, whereas class 4 cannot be identified. Therefore, the 4,645 subtomograms in class 1, 2, and 3 are likely to be true-positives missed by the template matching approach. For comparison, there are 2,843 ribosomes uniquely detected by the template matching approach. Since this number if about half of the 6,768 ribosomes uniquely detected by DISCA, we classified them into 2 classes using the same *RELION* procedure. The results shown in **Fig. S8** confirmed that they are truly ribosomes. Therefore, we empirically determined that DISCA has a false-positive rate of 9.3% and a false-negative rate of 15.0% (3.1% due to the particle picking preprocessing step). Moreover, DISCA detected about 20% of ribosomes missed by the template matching approach. There are 23,592 true ribosomes detected by DISCA and template matching in total, which corresponds to our estimated number of all ribosomes in these 65 *Mycoplasma pneumoniae* cellular tomograms. Overall, DISCA detected more true ribosomes than template matching (20,749 vs 18,987).

**Figure S8:**
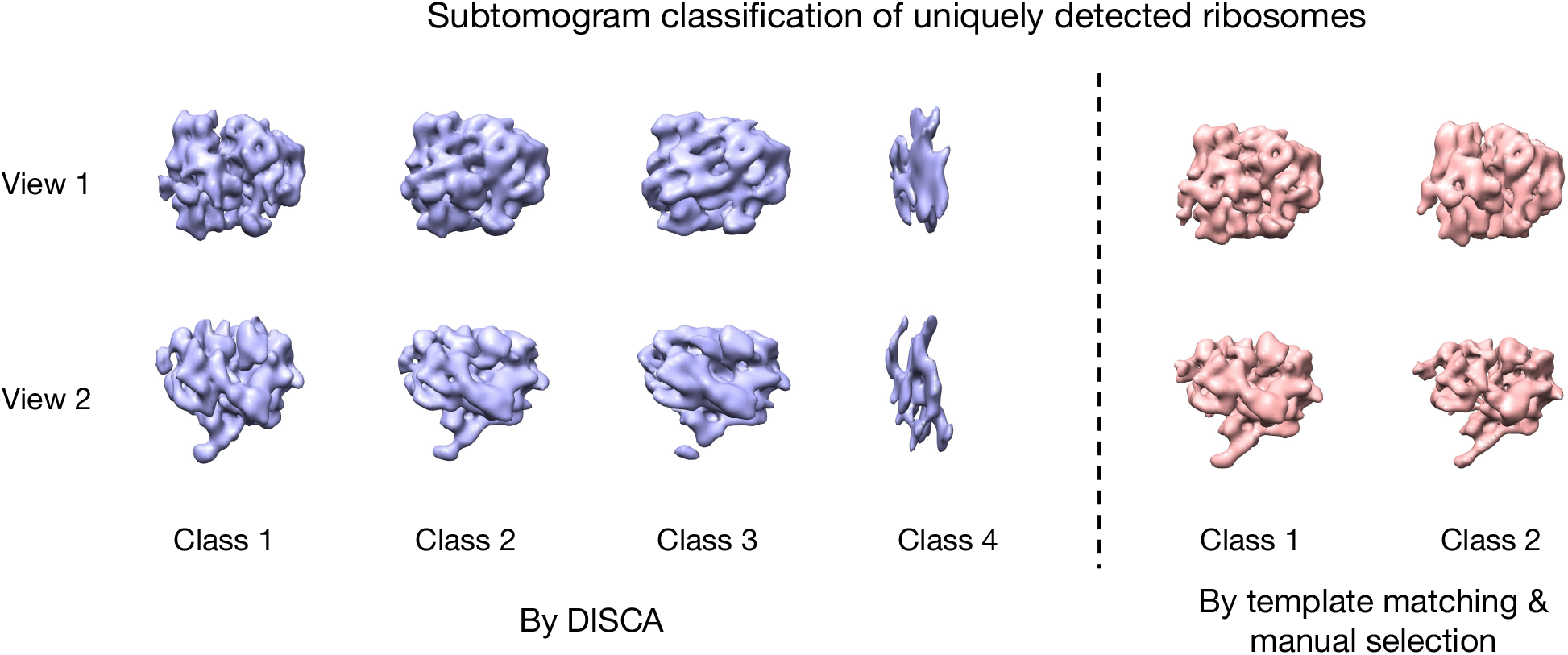
*RELION* subtomogram classification of uniquely detected ribosomes by the two approaches.

**Fig. S9** shows the comparison of example raw tomogram slice and corresponding re-embedding annotation of discovered patterns from a set of seven *Rattus* neuron tomograms [3]. The identified clusters consist of 12,229 subtomograms from 38,292 total extracted subtomograms. The voxel size of this tomogram is 13.68 Å and resolution measured on the ribosome pattern (averaged from 3,708 subtomograms) is 27.36 Å. The original article (Figure 2 in [3]) recovered three macromolecular complexes by manual curation, template matching, and subtomogram classification: ribosome, proteasome, and TRiC, which were validated by our results.

**Figure S9:**
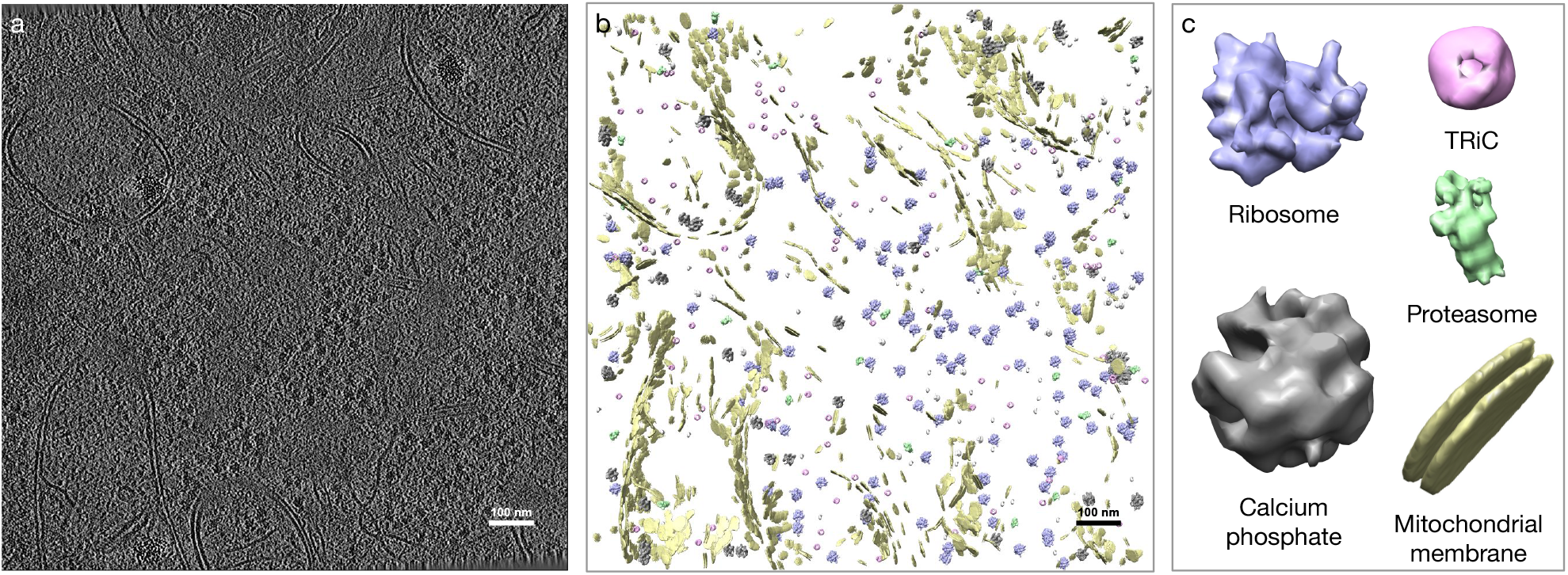
Example unsupervised annotation on a *Rattus* neuron tomogram [3]: a. slice of the original tomogram. b. discovered patterns re-embedded to the original tomogram space. c. iso-surface visualization of discovered patterns identified (generated from subtomogram averaging).

**Fig. S10** shows the comparison of example raw tomogram slice and corresponding re-embedding annotation of discovered patterns from a set of two *Synechocystis* cell tomograms [7]. The identified clusters consist of 4,804 subtomograms from 12,912 total extracted subtomograms of voxel size 13.68 Å. Since this is a small dataset with two tomograms, the ribosome pattern (averaged from 680 subtomograms) is not as ideal as other datasets. Nevertheless, DISCA validated the ribosome and thylakoid-attached phycobilisome array structures recovered by manual curation and subtomogram classification as in the original article [7].

**Figure S10:**
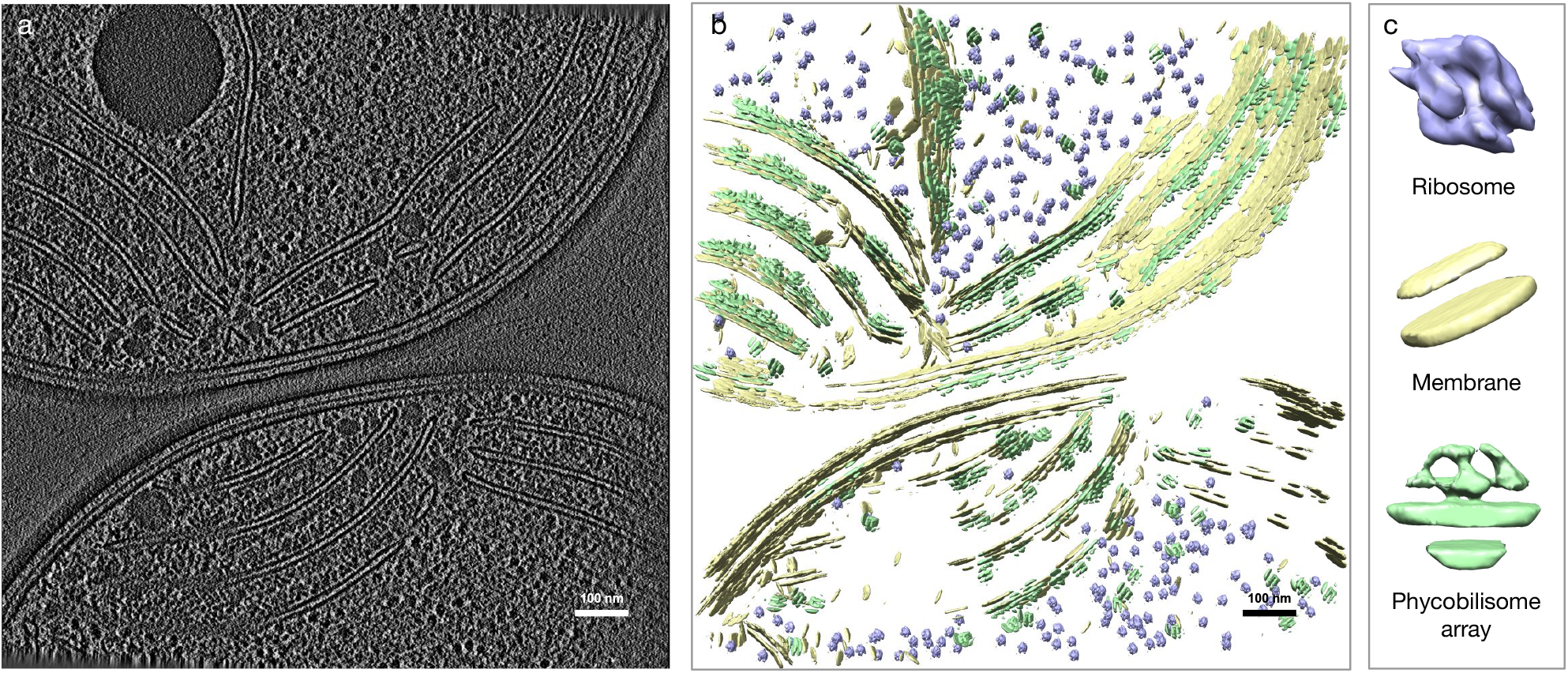
Example unsupervised annotation on a *Synechocystis* cell tomogram [7]: a. slice of the original tomogram. b. discovered patterns re-embedded to the original tomogram space. c. iso-surface visualization of discovered patterns identified (generated from subtomogram averaging).

**Fig. S11** compares example raw tomogram slices and corresponding re-embedding annotations of discovered patterns from a set of two *Cercopithecus aethiops* kidney cell tomograms [5]. The identified clusters consist of 2,219 subtomograms from 10,444 total extracted subtomograms of voxel size 14.20 Å. Since this is a small dataset with two tomograms, the ribosome pattern (averaged from 1,277 subtomograms) is not as ideal as other datasets. The original article [5] reported coarse discovery of globular and surface pattern using an autoencoder clustering model on this dataset. However, the ribosome-like globular pattern is of very low resolution, which is probably due to the impurity of the resulting clusters. DISCA showed notable improvement on this dataset as compared to Fig. 11 and S5 of the original article [5].

**Figure S11:**
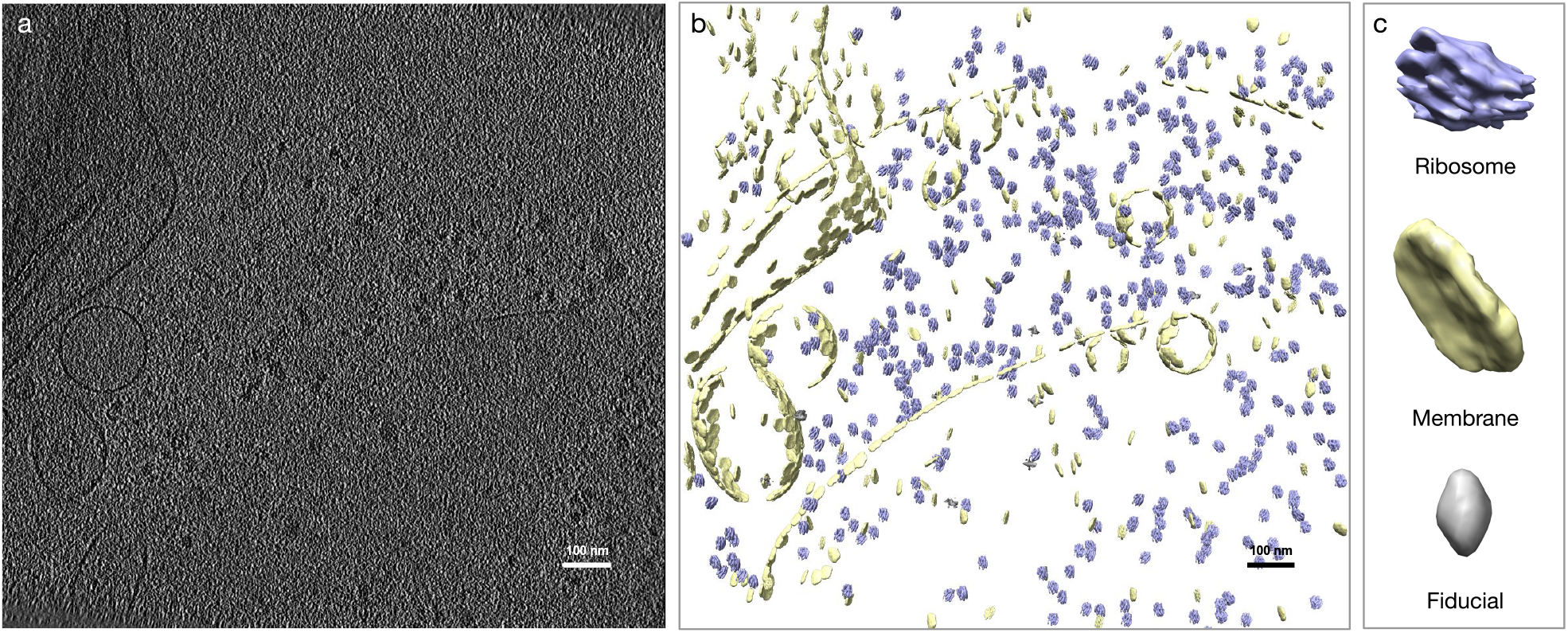
Example unsupervised annotation on a *Cercopithecus aethiops* kidney cell tomogram [5]: a. slice of the original tomogram. b. discovered patterns re-embedded to the original tomogram space. c. iso-surface visualization of discovered patterns identified (generated from subtomogram averaging).

**Fig. S12** compares example raw tomogram slices and corresponding re-embedding annotations of discovered patterns from a set of twenty *Murinae* embryonic fibroblast tomograms obtained from ETDB [11]. The identified clusters consist of 11,471 subtomograms from 54,684 total extracted subtomograms. The voxel size of this tomogram is 15.48 Å and resolution measured on the ribosome pattern is 33.77 Å (averaged from 2,459 subtomograms).

**Figure S12:**
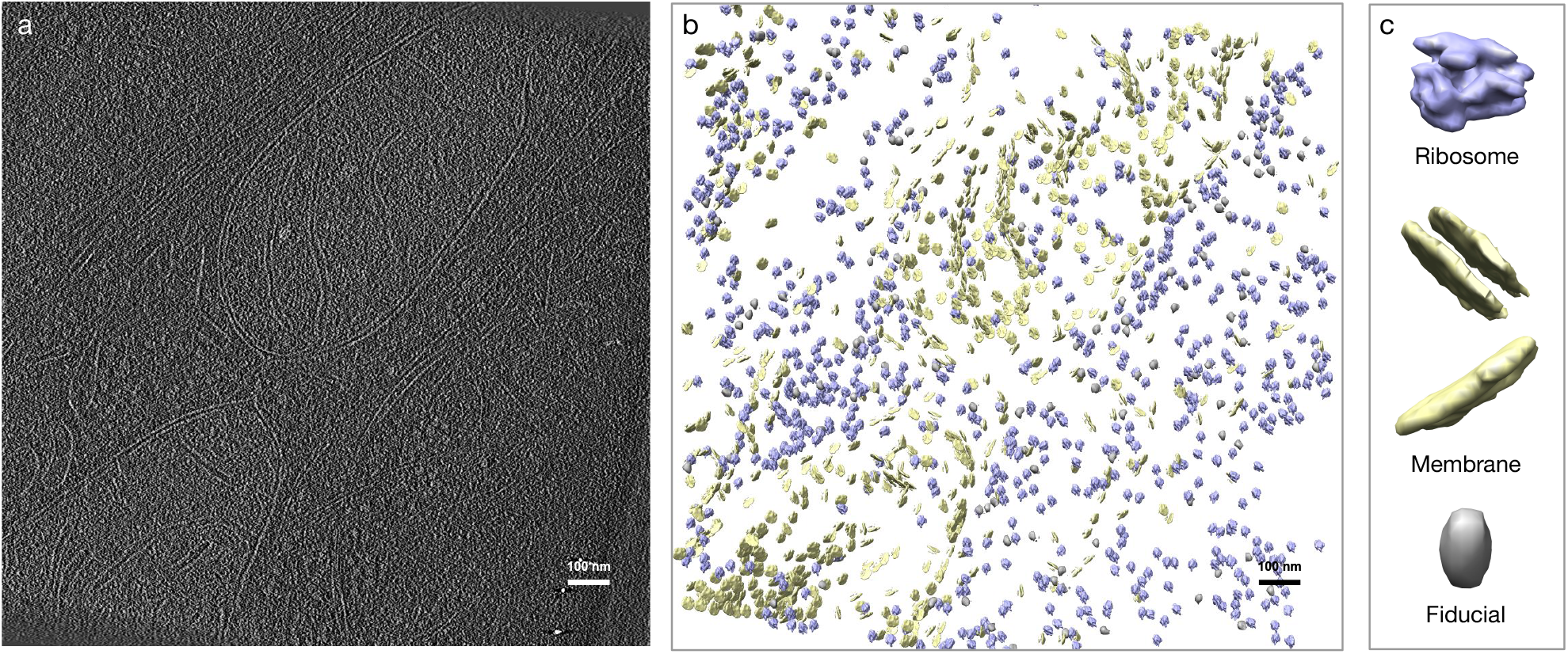
Example unsupervised annotation on a *Murinae* embryonic fibroblast cell tomogram [11]: a. slice of the original tomogram. b. discovered patterns re-embedded to the original tomogram space. c. iso-surface visualization of discovered patterns identified (generated from subtomogram averaging).

### Supplementary note 3: Fast visualization of cluster centers through a decoder

To validate that the learned features encode essential structural information of the input subtomograms, we trained a decoder using the *Rattus* neuron cryo-ET dataset [3] as an example. The input to the decoder is the learned features from DISCA and the output is the reconstruction of the input 3D subtomograms. Similar to [5], we then decoded the cluster centers, arithmetic averages of all feature vectors in a cluster, into reconstructed 3D images. Alternatively, instead of cluster center, features closest to each cluster center can also be decoded, which yield similar results. As shown in **Fig. S13**, center decodings of identifiable clusters resemble the type of structures contained, which validates the essential structural information effectively learned by the extracted features. Center decodings of non-identifiable clusters mostly resemble a tiny globular structure, which likely to indicates that most subtomograms contained in these clusters are either noises or structures too small. Therefore, DISCA can be used to efficiently filter out false-positive particles picked by template-free particle picking methods.

**Figure S13:**
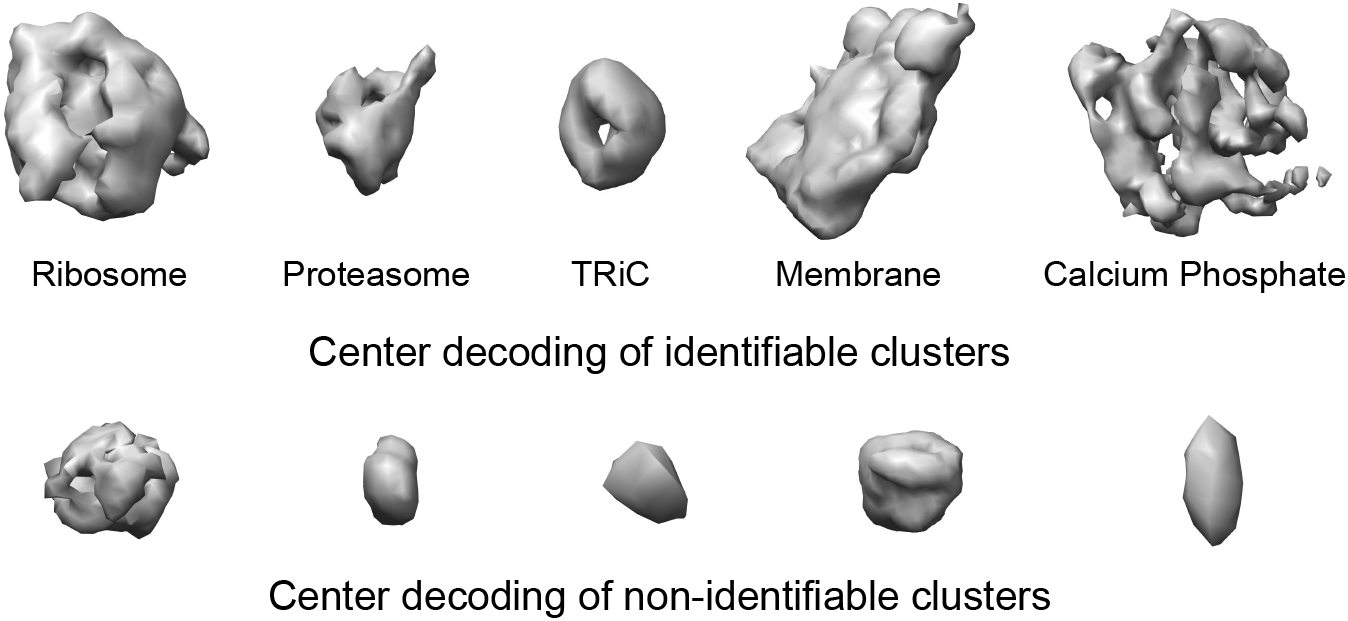
Example decodings of cluster centers from the *Rattus* neuron dataset.

In addition, it is very useful to quickly identify interesting clusters for downstream analysis before doing the computationally intensive subtomogram averaging step. The training of the decoder from scratch on this dataset of 36,377 subtomograms took less than 10 minutes. Therefore, the decoding of cluster centers can be used for such identification purpose, especially for structural clusters that can be easily recognized such as ribosome, surface patterns, and fiducial markers.

**Figure S14:**
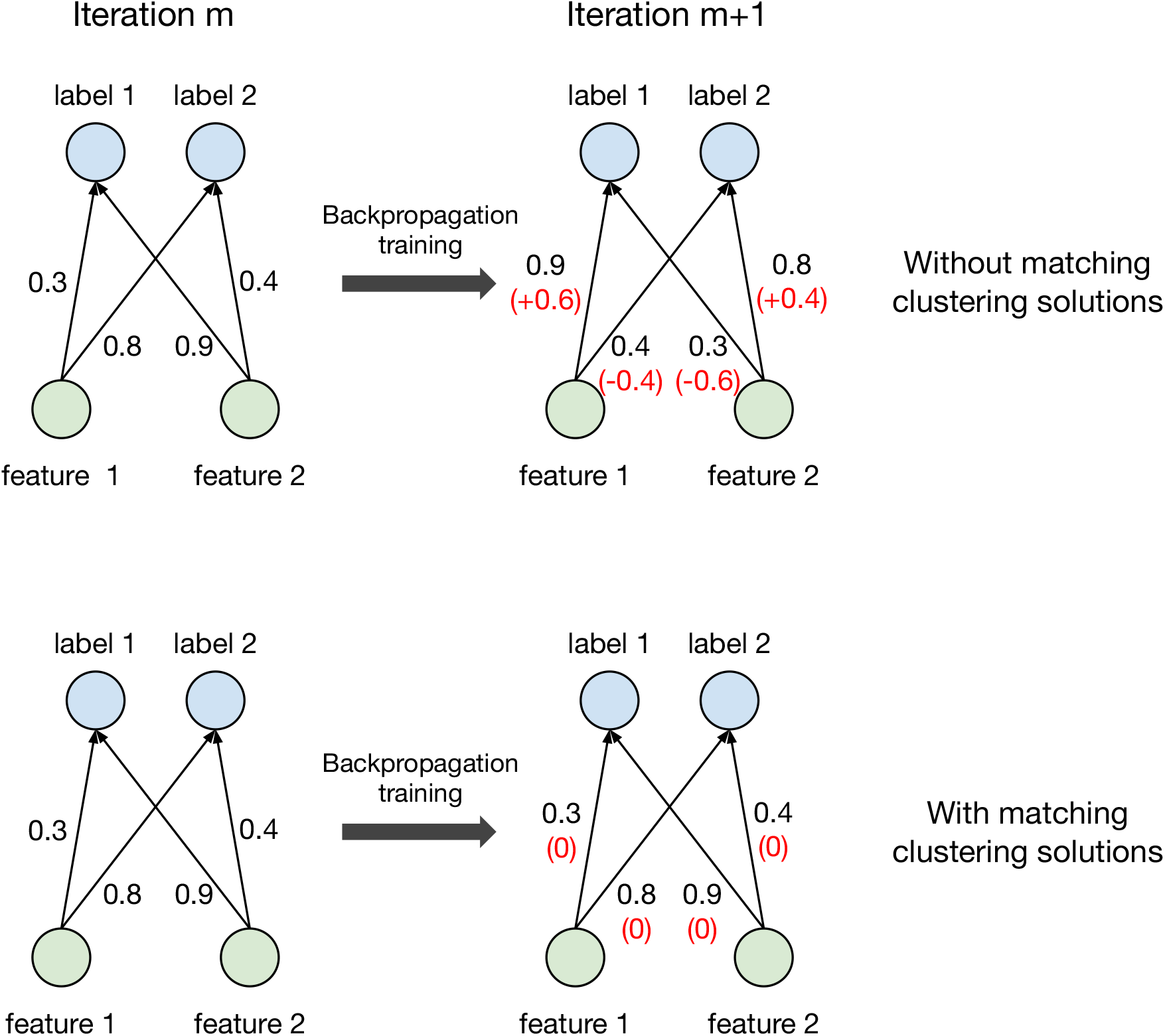
We assume the last fully connected layer for classification has two input feature nodes and two output label nodes. And we assume the clustering solution has two clusters 1 and 2 with labels flipped from iteration *m* to iteration *m*+1. Without matching clustering solutions, the backpropagation training needs to re-learn (large changes in weights) the already optimized weights to correctly output the flipped labels. This will cause strong instability during training. However, with matching clustering solutions, the already optimized weights no longer need to be re-learned (no change in weights).

### Supplementary note 4: Implementation details

The neural network model YOPO was implemented in platform Keras [12] with Tensorflow backend [13]. No external pre-trained models or additional supervision were used. All models were trained on a computer with 4 NVIDIA GeForce Titan X Pascal GPU instances and 48 CPU cores. The statistical model fitting used functions in Python package *numpy* and *sklearn*. The implementation of the Hungarian algorithm used functions in Python package *scipy*. The data augmentation used random 3D rotation functions implemented in *AITom* [14]. During the YOPO model training, the label smoothing factor gradually decreases by a factor of 0.9 in each iteration as we expect the amount of mislabeled data to decrease over time, and therefore YOPO becomes more certain about its prediction over time. During the Gaussian mixture model fitting, the extracted features are dimension reduced by PCA to length of 32 as an optional step for faster clustering. To measure the convergence of DISCA, a generalized EM framework, we set two convergence criteria: (1) the estimated *K* and the vast majority (99%) of the estimated labels stay the same for three consecutive iterations, or (2) the maximum number of iterations has been reached.

#### Preprocessing

For template-free particle picking, we applied the 3D Difference of Gaussians (DoG) [15] volume transform algorithm implemented in *AITom*. 3D DoG first computes a map *I*_*DoG*_ by subtracting two Gaussian blurred versions of the input tomogram *v* using the Gaussian function *I* with different standard deviations *σ*_1_ and *σ*_2_, where, without loss of generality, *σ*_1_ *> σ*_2_. The 3D DoG map is computed on tomogram *v* as *I*_*DoG*_ = *I*_*v*_(*σ*_1_) −*I*_*v*_(*σ*_2_).

Local maxima are detected to extract a set of subtomograms *S* from *v* as:

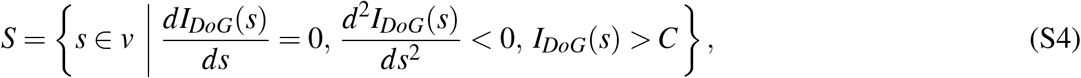

where *s* is a 3D location in *I*_*DoG*_ and *C* is a threshold applied for selecting local peaks. In our implementation, we ensured a minimum distance of 15 voxels between two peaks by filtering out peaks with low values.

#### Postprocessing

For subtomograms in each structurally homogeneous subset obtained from DISCA, iterative 3D averaging was performed using *RELION 3*.*0*. As a template-and-label-free framework, we did not use any external structural templates in the averaging process. The initial averages were obtained by our unsupervised deep learning based subtomogram alignment method Gum-Net (implemented in AITom) [14,16]. After the iterative 3D averaging process, the subtomogram averages were reembedded into the original tomogram by Gum-Net for visualization purposes.

